# Ligand binding promiscuity of αVβ3 integrin is enlarged in response to mechanical force

**DOI:** 10.1101/200493

**Authors:** Michael Bachmann, Markus Schäfer, Vasyl V. Mykuliak, Marta Ripamonti, Lia Heiser, Kai Weißenbruch, Sarah Krübel, Clemens M. Franz, Vesa P. Hytönen, Bernhard Wehrle-Haller, Martin Bastmeyer

## Abstract

αVβ3 integrin recognizes multiple extracellular matrix proteins, including vitronectin (Vn) and fibronectin (Fn). However, cell experiments are frequently performed on homogenously coated substrates with only one integrin ligand present. Here, we employed binary-choice substrates of Fn and Vn to dissect αVβ3 integrin-mediated binding to both ligands on the subcellular scale. Superresolution imaging revealed that αVβ3 integrin preferred binding to Vn under various conditions. In contrast, binding to Fn required mechanical load on αVβ3 integrin. Integrin mutations, structural analysis, and molecular dynamics simulations established a model where the extended-closed conformation of αVβ3 integrin binds Vn but not Fn. Force-mediated hybrid domain swing-out characterizes the extended-open conformation needed for efficient Fn binding. Thus, force-dependent conformational changes in αVβ3 integrin increase the number of available ligands and therefore the ligand promiscuity of this integrin. These findings for αVβ3 integrin were shown to regulate cell migration and mechanotransduction differentially on Fn compared to Vn and therefore to regulate cell behavior.

## Introduction

Integrins, an important class of cell-matrix adhesion receptors, consist of α- and β-subunits forming transmembrane heterodimers (Campbell and Humphries, 2011). While the extracellular part binds to proteins of the extracellular matrix (ECM), the intracellular part is connected to actin via multiple adapter and signaling proteins that make up the so-called adhesome (Hytönen and Wehrle-Haller, 2014, Zaidel-Bar et al., 2007). The functional definition of integrin activation is based on the capacity to bind ligands. The structural definition associates the active structure with conformational changes from a bent, to an extended-closed, and finally to an extended-open state (Campbell and Humphries, 2011, Zhu et al., 2008, Su et al., 2016). Concerning the β-integrin subunit, the extended-open conformation is characterized by the swing-out of the hybrid domain in respect to the βA domain (Zhu et al., 2013). Molecular dynamics (MD) simulations indicate that force supports this hybrid domain swing-out and thereby leads to full integrin activation (Zhu et al., 2008, Puklin-Faucher et al., 2006, Dong et al., 2017). These theoretical studies are supported by multiple examples where mechanical forces regulate integrin functions, structural elements of the adhesome, and thus integrin-mediated adhesions (Balaban et al., 2001, Stricker et al., 2011, Hytonen and Wehrle-Haller, 2016, Nordenfelt et al., 2016, Rahikainen et al., 2017, Kuo et al., 2011, Schiller et al., 2011).

The best-studied class of integrins is the group of RGD-binding integrins sharing the ability to interact with an exposed Arg-Gly-Asp peptide in their ligands (Hynes, 2002). Promiscuity between ligands and receptors is especially common for these RGD-integrins. For example, αVβ3 integrin is reported to have the ability to recognize at least 12 different ligands (Humphries et al., 2006). Initially described as the vitronectin (Vn) receptor (Pytela et al., 1985), αVβ3 integrin is now widely accepted and analyzed in its function as a fibronectin (Fn) receptor (Roca-Cusachs et al., 2009, Schiller et al., 2013, Cao et al., 2017, Danen et al., 2002, Balcioglu et al., 2015). However, in these studies αVβ3 integrin was challenged with one ligand at a time. The ECM, in contrast, is more complex and might offer different ligands in close distance to each other potentially changing integrin behavior dependent on the ligand.

To address this issue of substrate heterogeneity, we have recently developed a method to produce microstructured, differential Fn/Vn substrates that enable us to analyze the influence of Fn and Vn coated surfaces on a subcellular scale (Pinon et al., 2014, Rahikainen et al., 2017). Here, we combined this method with super-resolution microscopy and super-resolution live cell imaging. We revealed a force-dependent ligand binding of αVβ3 integrin to Fn, while this integrin binds to Vn already under low mechanical load. Further experiments explain this behavior based on different integrin conformations. Finally, we found that Vn, together with osteopontin (Opn), belongs to a class of high-affinity ligands for αVβ3 integrin contrasting with the low-affinity, force-induced ligands Fn and fibrinogen (Fbg).

## Results

### Vitronectin is the preferred ligand for αVβ3 integrin

To study how the simultaneous presentation of two ECM ligands influences binding of αVβ3 integrin, we produced binary Fn/Vn substrates with subcellular structural resolution (Pinon et al., 2014). 2×2 μm squares of Fn separated by 1 μm gaps were stamped onto a coverslip and the remaining surface was covered with Vn leading to a geometrical coverage of about equal contribution (Pinon et al., 2014). The quality of substrates was analyzed by fluorescence and atomic force microscopy (Fig. 1a, b). To reveal integrin specific ECM ligand binding, β3-wt integrin GFP was transfected into NIH 3T3 cells expressing low levels of endogenous β3 integrins (Pinon et al., 2014), and cultured on the patterned surfaces for 2 h. In these cells, αV integrin is the only subunit pairing with β3 integrin. Thus, our results obtained with β3-wt integrin GFP are synonymous for αVβ3 integrin. Subsequent fixation and immunostaining for paxillin allowed detecting all integrin-mediated cell-matrix adhesions. Imaging with super-resolution structured illumination microscopy (SR-SIM) revealed extended overlap of αVβ3 integrin and anti-paxillin staining (Fig. 1c). However, some paxillin-clusters on Fn were free of αVβ3 integrin, indicating recruitment to endogenous Fn-bound α5β1 integrin (Supplementary Fig. 1b, g). Therefore, these substrates enabled us to analyze the influence of Fn and Vn on αVβ3 integrin adhesion formation at a micrometer scale within single cells. Surprisingly, we found a clear preference of αVβ3 integrin-positive adhesions for Vn with 83.5 % colocalization (Fig. 1d). This preference for Vn was independent of the size of the αVβ3 integrin-mediated adhesion (Fig. 1e), the presence of α5β1 integrin (Supplementary Fig. 1a, f), or the stamping procedure (Supplementary Fig. 1d, j). Moreover, combining Fn/Vn patterns with hydrogels confirmed this preference of αVβ3 integrin for Vn over a wide stiffness range (Supplementary Fig. 1k-m). Homogenous substrates supported these findings showing about twofold reduced αVβ3 integrin-mediated adhesion area and recruitment into adhesions on Fn compared to Vn (Supplementary Fig. 2).

**Figure 1:**
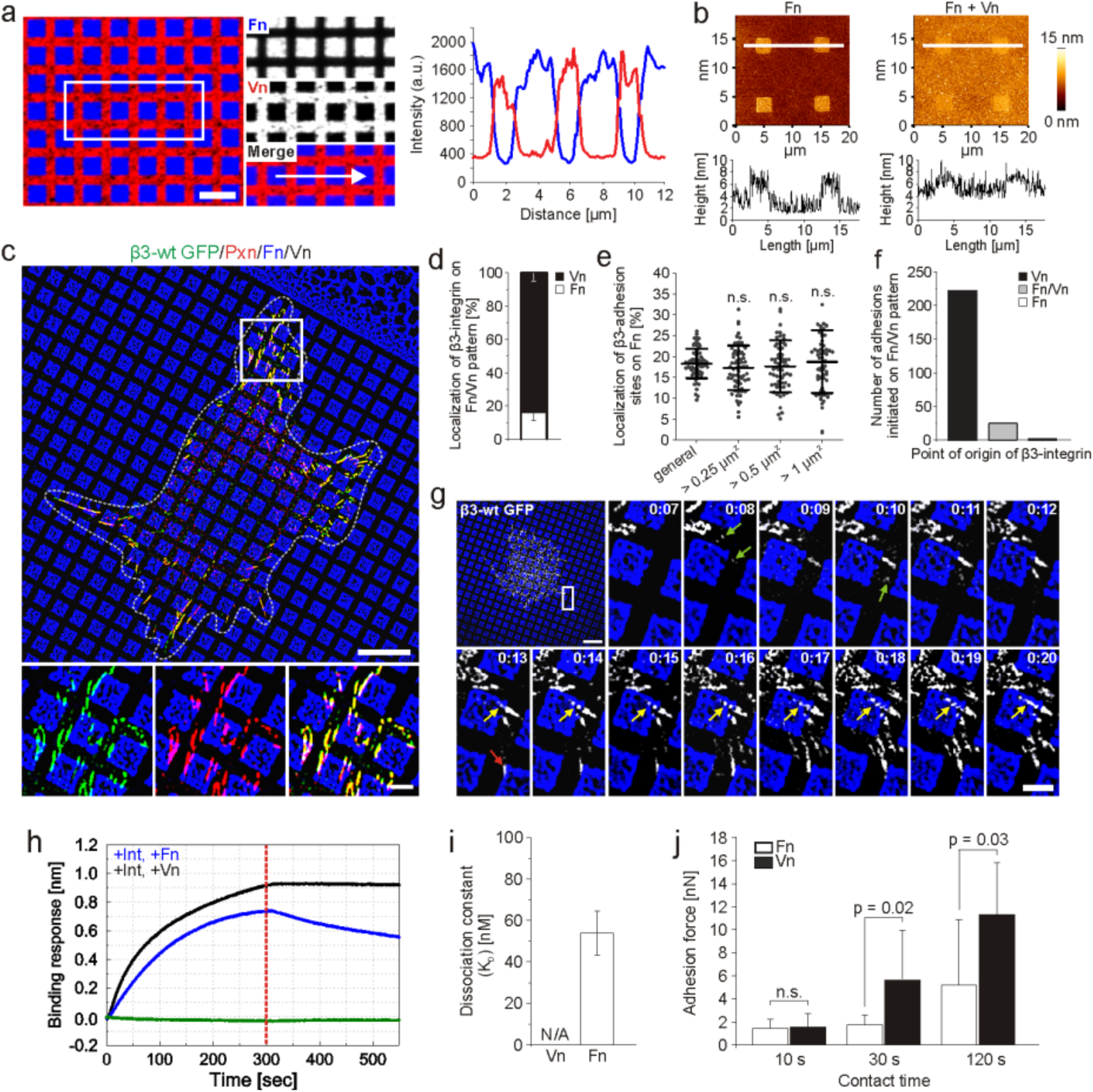
αVβ3 integrin favors binding to vitronectin (Vn) compared to fibronectin (Fn). (**a**) Microcontact printing of 2×2 μm squares of Alexa Fluor 647 labeled Fn (blue) onto glass cover slips and backfilling the pattern with Alexa Fluor 568 labeled Vn (red) leads to differential Fn/Vn patterns as shown by the fluorescence intensity profile along the arrow. Geometrical coverage varies dependent on the pattern quality: 44-49% Fn, 56-51% Vn. (**b**) Height profiles of Fn patterns (left) and Fn/Vn patterns (right) measured with atomic force microscopy (AFM) in contact mode. Profiles along the white lines indicate a monolayer of Fn and a uniform topography of the binary choice substrates. (**c**) Super-resolution structured illumination microscopy (SR-SIM) image of a NIH 3T3 cell transfected with β3-wt GFP integrin (green), cultured on a Fn/Vn pattern (Fn in blue), and immunostained for paxillin (red). The cell contour is outlined with a dashed white line. (**d**) Quantification of colocalization of β3-wt GFP integrin with Fn or Vn for fixed cells (n = 66; N = 4). (**e**) Quantification of colocalization of β3-wt GFP integrin with Fn for all adhesions (“general”) or only for those bigger than the indicated threshold (re-analysis of the data from d). (**f**) Total number of αVβ3-mediated adhesions that initiated on Vn, Fn, or at the Fn/Vn-interface for cells imaged with live cell SR-SIM. Quantification is based on 246 initiated adhesions from six cells out of three independent experiments. (**g**) NIH 3T3 cell transfected with β3-wt GFP integrin (white) seeded on Fn/Vn patterns (Fn in blue) was monitored with live cell SR-SIM (also see Supplementary Video 1). Magnifications show initiation and maturation of αVβ3-mediated adhesions (time is given in h:min). Green arrows point to newly established adhesions. Yellow arrows follow adhesions that initiated on Vn while they translocate to Fn. The red arrow at 13 min indicates an adhesion that appeared at the Fn/Vn-interface. (**h**) Representative curves for Fn and Vn association (0 sec to 300 sec) and dissociation (300 sec to 550 sec) to αVβ3 integrin measured with biolayer interferometry. (**i**) K_D_ values for the interaction of purified αVβ3 integrin to Vn or Fn measured with biolayer interferometry (values calculated from n = 9 measurements in N = 3 independent experiments). (**j**) Single-cell force spectroscopy of GD25 cells (expressing αVβ3, but no β1 integrin). Detachment forces on Fn and Vn measured after the indicated contact time points. Typically 10 force measurement repetitions were performed for each cell and time point, and a total of 8 cells were tested. (**a, c, g**) Scale bars: 10 μm in overviews, 2 μm in zoom-ins.

To study the dynamics of αVβ3 integrin-mediated adhesion formation, we applied SR-SIM live cell imaging. During spreading, cells initiated numerous nascent adhesions (Fig. 1g and Supplementary Video 1 and 2). These adhesions almost exclusively appeared on Vn (Fig. 1f) while some of them were later translocated towards Fn in a centripetal direction towards the cell center (yellow arrows in Fig. 1g and Supplementary Video 2). In control experiments where Fn was replaced by albumin, αVβ3 integrin-mediated adhesions remained restricted to Vn (Supplementary Fig. 1c), indicating that ligand binding was necessary for localization and stabilization of αVβ3 integrins on Fn-coated squares. Next, we asked whether the preference for Vn could be explained by different affinities of αVβ3 integrin for these two ligands. Using biolayer interferometry, we analyzed the dissociation constant of purified αVβ3 integrins for Fn and Vn (Fig. 1h, i). Interestingly, Vn showed measurable dissociation from αVβ3 integrins only in the presence of the αVβ3 integrin inhibitor cilengitide (w/o cilengitide: K_D_ < 1 pM; + 20 μM cilengitide: K_D_ = 302.6 nM; Supplementary Fig. 3i-k), which was similar to previous reports showing non-dissociable binding to Vn (Orlando and Cheresh, 1991). Fn, in contrast, dissociated rapidly (K_D_ = 53.98 nM), proposing that the higher affinity of the extracellular domain of αVβ3 integrin for Vn explains the observed Vn-preference. Accordingly, a chimeric integrin with extracellular β3 and intracellular β1 domains (β3/β1 chimera) showed the same preference for Vn as β3-wt integrin on the micropatterned Fn/Vn surfaces (Supplementary Fig. 1e, h). In addition, we performed atomic force microscopy (AFM)-based single-cell force spectroscopy (Langhe et al., 2016, Dao et al., 2012) on substrates with adjacent Fn and Vn areas. A living cell attached to the AFM cantilever was alternatingly brought in contact with and retracted from two homogenously coated but adjacent Fn and Vn areas. With respect to different contact times (typically ten force cycles per cell and contact time), the detachment force was determined from force-distance curves collected during cell retraction. Starting after 30 s of adhesion time, a significantly higher force was needed to detach cells from Vn compared to Fn substrates (Fig. 1j).

Together, our experiments revealed that initial αVβ3 integrin-mediated adhesions formed exclusively on Vn, potentially explained by the higher binding affinity of αVβ3 integrin to Vn. However, the question remained why the colocalization of αVβ3 integrin-mediated adhesions with Fn was specific for maturing focal adhesions.

#### Actomyosin contractility regulates the ligand preference of αVβ3 integrin

αVβ3 integrin-mediated adhesions translocated onto Fn squares strictly towards the cell center in the direction of retrograde actin flow. To test whether actomyosin forces are involved in the Fn/Vn selection process of αVβ3 integrin, we reduced actomyosin contractility with blebbistatin or Y27632. The latter compound inhibits the Rho/ROCK pathway and thereby reduces myosin activity, while blebbistatin inhibits myosins directly. As previously shown (Choi et al., 2008), both inhibitors increased the number of small, round nascent adhesions in the cell periphery (Fig. 2a, b). In addition, however, treatment with both inhibitors led to a significant decrease in colocalization of β3-wt GFP integrin with Fn (Fig. 2g and Supplementary Fig. 3a). Culturing β3-wt GFP expressing cells on Fn/BSA or Vn/BSA binary choice substrates confirmed that reduction of contractility affected αVβ3 integrin more on Fn (Supplementary Fig. 3e, f) compared to Vn (Supplementary Fig. 3g, h). Vinculin is well established as an important part of the molecular clutch to transmit forces from actin to the integrin-ECM bond (Humphries et al., 2007, Thievessen et al., 2013, Rahikainen et al., 2017). Therefore, we analyzed the localization of β3-wt GFP integrin on Fn/Vn substrates for mouse embryonic fibroblasts derived from vinculin knockout mice (MEF Vcl -/-). Indeed, the absence of vinculin led to a decrease in Fn binding of αVβ3 GFP integrin (Fig. 2e, h) that was comparable to the results obtained with the contractility inhibitors (Fig. 2g). Control experiments using wt MEFs (Fig. 2d) and Vcl -/- MEFs re-expressing vinculin mCherry (Fig. 2f) demonstrated that the localization of αVβ3 GFP integrin on Fn can be rescued by vinculin expression (Fig. 2h). To increase the mechanical load on integrins, we overexpressed non-muscle myosin IIA mApple (NMIIA) in NIH3T3 cells (Fig. 2c). Adding additional NMIIA caused a significant increase of αVβ3 GFP integrin localization on Fn (Fig. 2g). Taken together, these findings indicate that αVβ3 integrin binds to Vn irrespective of the mechanical load whereas Fn binding is fostered by intracellular force and vinculin.

**Figure 2:**
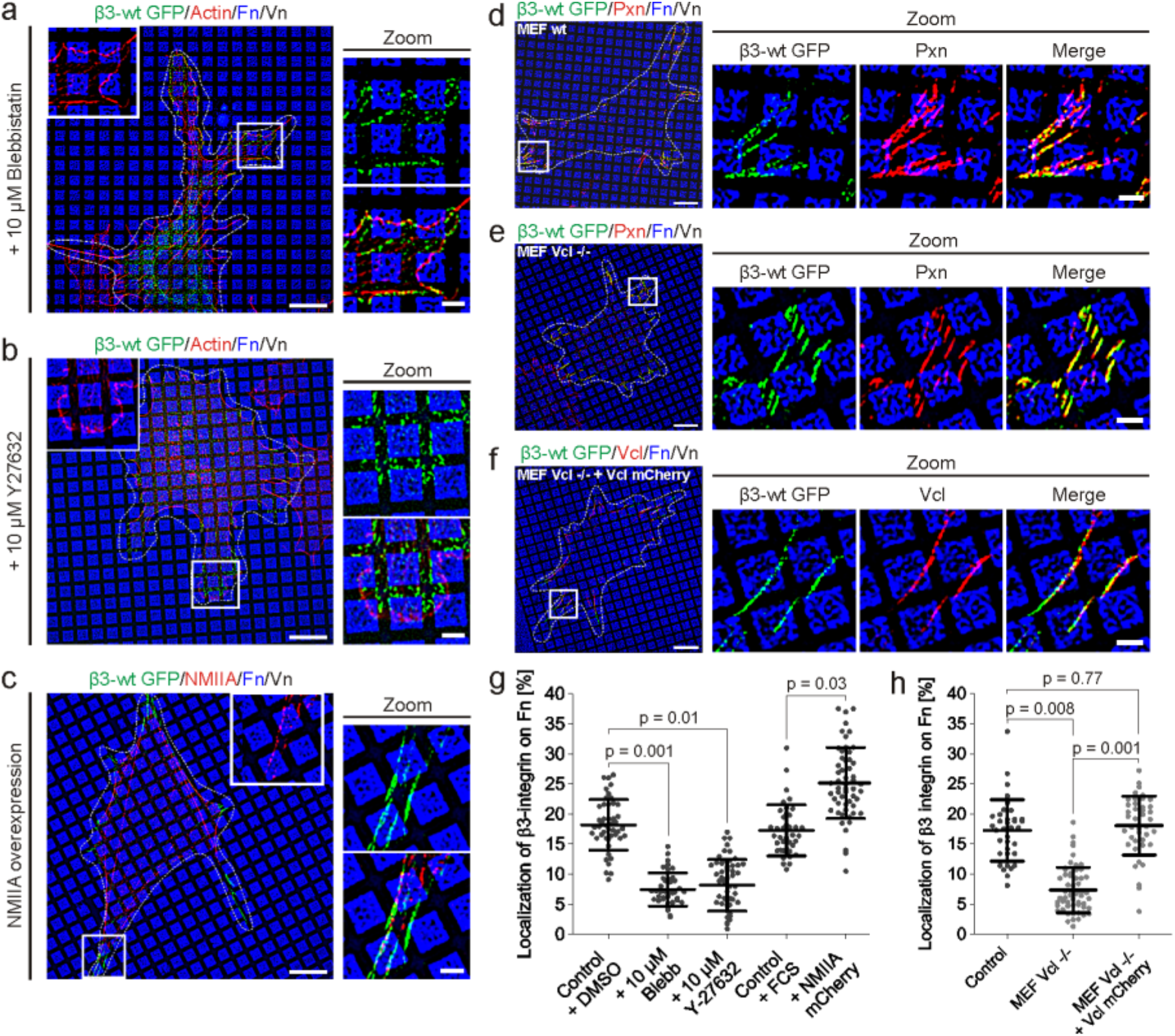
Mechanical force regulates ligand preference of ¤Vß3 integrin. (**a**) NIH 3T3 cells transfected with β3-wt GFP integrin (green) were cultured on Fn/Vn substrates (Fn in blue) in the presence of blebbistatin and were stained after fixation for actin (red). (**b**) NIH 3T3 cells treated with Y27632. (**c**) NIH 3T3 cells transfected with β3-wt GFP integrin (green) and myosin IIA mApple (NMIIA; red) with serum (FCS) present in the medium. (**d**) MEF wt or (**e**) MEF vinculin knockout cells (MEF Vcl -/-) transfected with β3-wt GFP integrin (in green) and immunostained for paxillin (Pxn; red). (**f**) MEF Vcl - /- cells transfected with β3-wt GFP integrin (green) and Vcl mCherry (red). (**g**) Quantifications of colocalization of β3-wt GFP with Fn for cells treated as described in **a-c** (control + FCS: n = 66, N = 3; control + DMSO: n = 46, N = 3; + 10 μM blebbistatin: n = 40, N = 3; + 10 μM Y-27632: n = 54, N = 3; + NMIIA mCherry: n = 55, N = 3). (**h**) Quantifications of colocalization of β3-wt GFP with Fn for cells treated as described in **d-f** (control: n = 38, N = 3; MEF Vcl -/-: n = 57, N = 3; MEF Vcl -/- + Vcl mCherry: n = 42, N = 3). (**a-f**) All fluorescent images were acquired with SR-SIM. White dashed lines indicate the cell outline. Scale bar: 10 μm in overview images, 2 μm in zoom ins.

#### Hybrid domain swing-out is required for Fn binding of αVβ3 integrin

Next, we asked whether increased activation of αVβ3 integrin might substitute force in the process of Fn binding. Therefore, we employed Mn^2+^ activation of αVβ3 integrin (Fig. 3a) or established integrin mutations (Fig. 3b-d): (i) Mn^2^+ treatment increases the affinity of the integrin headpiece for the ligand (Zhu et al., 2013), (ii) the β3-VE mutant increases the affinity for talin 20-fold leading to increased integrin activation (Pinon et al., 2014), (iii) β3-D723A disrupts the inhibitory salt-bridge at the inner membrane clasp between the αV- and β3-subunits (Saltel et al., 2009), and (iv) β3-N305T is reported to cause a constitutive hybrid domain swing-out and slower integrin dynamics (Luo et al., 2003, Cluzel et al., 2005). Surprisingly, on Fn/Vn binary choice substrates, only β3-N305T showed a significant increase of colocalization with Fn (Fig. 3f), while Mn^2^+ treatment and the intracellular activating mutations (β3-VE and β3-D723A) caused no significant difference. Next, we tested whether the conformational changes caused by the β3-N305T mutation are accompanied by the ability to initiate adhesions on Fn by using live cell SR-SIM (Fig. 3j and Supplementary Video 3, 4) and compared this to the β3-VE mutation (Supplementary Video 5, 6). In both cases, spreading cells initiated most αVβ3 integrin-mediated adhesions on Vn. However, cells expressing β3-N305T were able to initiate numerous adhesions on Fn as well (Fig. 3k).

**Figure 3:**
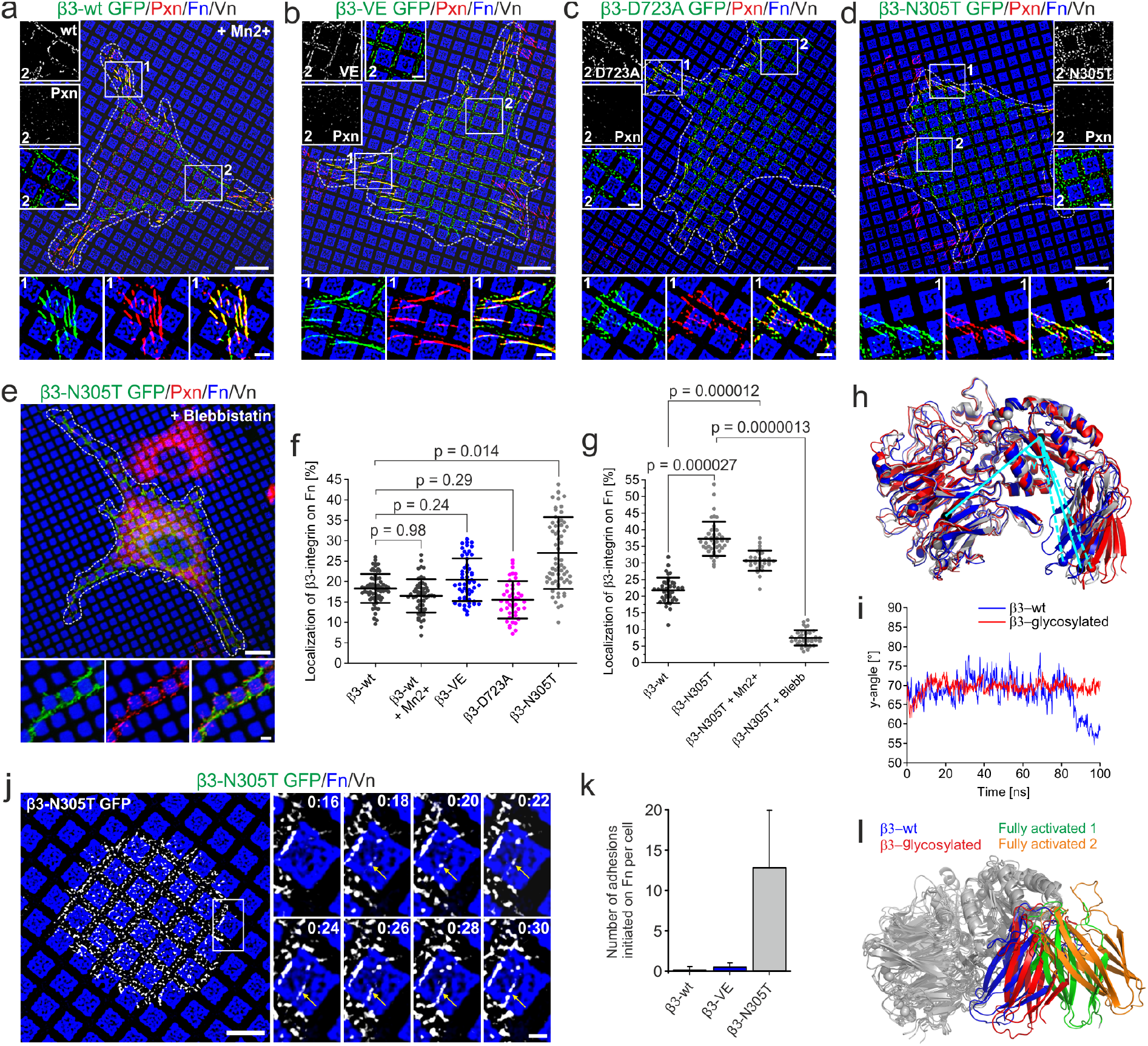
Hybrid domain swing-out is necessary for αVβ3 integrin binding to FN. (**a**) NIH 3T3 cell transfected with β3-wt GFP integrin (green). 1 mM Mn^2^+ was added to the medium 30 min before fixation. (**b**) NIH 3T3 cell transfected with β3-VE GFP integrin (green), or (**c**) for β3-D723A GFP integrin (green), or (**d**) for β3-N305T GFP integrin (green). (**e**) β3-N305T GFP integrin transfected cells treated with 10 μM blebbistatin. All cells (**a-e**) were fixed and immunostained for paxillin (Pxn; red). Zoom-ins depict adhesions in the cellular periphery (1) or the cell center (2). (**f**) Quantifications of colocalization of β3 GFP with Fn for cells treated as described in a-d and imaged with SR-SIM. Paxillin was used as a mask to exclude nascent αVβ3 integrin clusters in the cell center from analysis (β3-wt: replot of the data from Fig. 1d; β3-wt + Mn2+: n = 55, N = 3; β3-VE: n = 54, N = 3; β3-D723A: n = 43, N = 3; β3-N305T: n = 66, N = 4). (**g**) Quantification as described in f for cells treated as described before. Data for ‘β3-N305T + Mn2+’ was acquired from cells treated as described in d but with addition of 1 mM Mn^2^+ for the last 30 min before fixation. All images were acquired with diffraction-limited microscopy (β3-wt: n = 40, N = 3; β3-N305T: n = 42, N = 3; β3-N305T + Blebb: n = 39, N = 3). (**h**) Superimposition of (grey) the initial structure of αVβ3 integrin, (blue) the same structure after 100 ns molecular dynamics (MD) simulation, and (red) the N303-glycosylated structure after 100 ns MD simulation. Cyan lines indicate the position of hybrid domain swing-out measurements. (**i**) Fluctuation of the angle γ between βA and hybrid domain over time during the MD simulation. (**j**) Live cell SR-SIM imaging of a NIH 3T3 cell transfected with β3-N305T GFP integrin (white) spreading on Fn/Vn substrates (Fn in blue). Yellow arrows indicate an αVβ3 integrin-mediated adhesion that initiated on Fn. (**k**) Number of αVβ3 integrin-mediated adhesions per cell that initiated on Fn for NIH 3T3 cells transfected with the indicated integrin (β3-wt is a replot of the data in Fig. 1f; β3-VE: six cells analyzed out of three independent experiments; β3-N305T: six cells analyzed out of four independent experiments). (**l**) Superimposition of αVβ3 integrin structures as described in (**h**) for β3-wt (blue) and β3-glycosylated (red). Fully activated structures were created based on PBD: 4MMX with an arranged hybrid domain swing-out according to (Fully activated 1, green) PBD: 3EZ2, or (Fully activated 2, orange) PBD: 3FCU. (**a-e, j**) Scale bars: overview images 10 μm, zoom-ins 2 μm. White dashed lines indicate the cell outline. Images were acquired with (**a-d, j**) SR-SIM or (**e**) diffraction limited microscopy.

Additionally, we made a peculiar observation with all activating conditions. They caused central clusters of αVβ3 integrin with irregular shapes compared to classical adhesions (Fig. 3 a-d, zoom-in 2). Interestingly, these clusters localized almost exclusively on Vn (Supplementary Fig. 3o). Analysis of integrin adapter proteins showed talin recruitment but no association of these αVβ3 integrin clusters with paxillin, vinculin, or actin stress fibers (Fig. 3a-d, Supplementary Fig. 3l-n). This indicates that they are not mechanically coupled and thus are under low intracellular force (compare to MEF Vcl -/- cells; Fig. 2e, h). Similar integrin clusters have been reported before to appear within minutes after Mn^2^+ addition (Cluzel et al., 2005, Saltel et al., 2009). The exclusive localization of these low-force αVβ3 integrins on Vn complements our experiments using contractility inhibitors and vinculin -/- cells and emphasizes the requirement of mechanical load on αVβ3 integrin to bind to Fn.

Combined, these experiments showed that only αVβ3 integrin activation by the N305T mutation increases Fn binding and allows initiating of adhesions on Fn. Surprisingly, other established activating conditions failed to initiate adhesion on Fn (β3-VE; Fig. 3k) and to increase Fn binding (β3-D732A, β3-VE, and Mn^2^+ treatment; Fig. 3f).

#### Complete force-dependent hybrid domain swing-out is necessary for Fn binding

The β3-N305T mutation leads to increased ligand-binding (Luo et al., 2003) and reduced integrin dynamics (Cluzel et al., 2005) which is attributed to the creation of a glycosylation site between the βA and the hybrid domain at Asn 303 (N303). This is proposed to cause a constitutive hybrid domain swing-out and thereby full integrin activation. The unique ability of β3-N305T to increase Fn binding (Fig. 3f) motivated us to study this mutation in more detail. Specifically, we set out to dissect the steric effect of N303-glycosylation in β3-N305T from force-induced hybrid domain swing-out (Zhu et al., 2008, Puklin-Faucher et al., 2006). To this end, we reduced actomyosin forces by adding blebbistatin to β3-N305T GFP transfected cells cultured on Fn/Vn substrates (Fig. 3e). β3-N305T mediated adhesions appeared less affected by blebbistatin compared to β3 wt forming many nascent adhesions under this condition (Fig. 2a). However, the colocalization of β3-N305T GFP with Fn was clearly reduced in the absence of force (Fig. 3g). We observed the same effect of reduced Fn-binding for all other activating conditions when combined with a blebbistatin treatment (Supplementary Fig. 3b). Thus, all integrin activating conditions relied on force for Fn binding by αVβ3 integrin. Even constitutive hybrid domain swing-out as reported for the β3-N305T mutation was not sufficient for efficient Fn binding. Seemingly, force caused additional conformational changes needed for Fn binding.

To understand the impact of force and glycosylation on αVβ3 integrin conformation we employed molecular dynamics (MD) simulations for a αVβ3 integrin structure that was glycosylated at N303. Zhu and colleagues published headpiece opening of αIIbβ3 integrin in eight steps (Zhu et al., 2013). We used a Fn-bound structure of αVβ3 integrin (PDB: 4MMX) and arranged a hybrid domain swing-out by superimposition with step seven (PDB: 3ZE1) in the activation cascade described by Zhu and colleagues. This structure was modified by adding a glycosylation at N303 and equilibrated for 100 ns. The same structure without glycosylation at N303 was used as a control. Our MD simulations showed that hybrid domains in both structures swing out to a similar angle with the glycosylated form appearing more stable (Fig. 3h, i). Accordingly, glycosylation at N303 stabilizes αVβ3 integrin in a conformation close to full activation. The final activation step (step 8 according to Zhu et al., 2013, PBD: 3ZE2), in contrast, is characterized by an increased hybrid domain swing-out (“Fully activated 1” in Fig. 3l). Another published structure of active β3 integrin (PBD: 3FCU) confirmed this correlation of full integrin activation and increased hybrid domain swing-out (“Fully activated 2” in Fig. 3l). Thus, comparison of the glycosylated to fully activated structures revealed that N303 glycosylation alone is not sufficient for full hybrid domain swing-out. Combining β3-N305T with Mn^2^+ slightly decreased Fn-binding (Fig. 3g) as it was the case for adding Mn^2^+ to β3-wt (Fig. 3f) potentially indicating that the effect of Mn^2^+ on αVβ3 integrin conformation is limited.

We conclude that our MD simulations, performed without mechanical load on the integrin, reflected the structure of β3-N305T integrin in experiments with contractility inhibition (Fig 3e, g). Thus, experiments and simulation imply that the final activation step with maximal hybrid domain swing-out stabilizes Fn binding and that this step is only achieved for αVβ3 integrin under mechanical load.

#### Vn binding is initiated for αVβ3 integrin in the extended-closed conformation

Experiments and MD simulations indicated that stable Fn-binding by αVβ3 integrin requires force-dependent hybrid domain swing-out. In contrast, αVβ3 integrin was able to bind Vn in experiments where cell contractility was reduced. Accordingly, it is tempting to speculate that αVβ3 integrin can bind Vn already in the extended-closed conformation. To test this, we set out to develop an integrin mutation that locks αVβ3 integrin in the extended-closed conformation. Therefore, we created a disulfide bridge between the βA and the hybrid domain to limit the degree of the hybrid domain swing-out (β3-V80C/D241C). Structural analysis supported our rationale for this mutation (Fig. 4a). We prepared a model, where cysteine mutations were introduced into extended-closed conformation of αVβ3 integrin (PDB: 4MMX) using PyMOL and energy minimization of the model. The disulphide bridge caused only minimal distortion for the protein; the distance between Cα atoms of V80C and D241C after introducing a disulphide bond did not change compared to the wildtype situation (both structures: d = 6.3 Ä). In contrast, αVβ3 integrin in the extended-open conformation showed an increased distance by a factor of three (d = 19.4 Ä) between Cα atoms of V80 and D241, indicating that a V80C-D241C disulphide bridge can block the transition to the extended-open conformation. The structure of the extended-open form was based on the closed integrin and opening was prepared by superimposition of βA and hybrid domains with the crystal structure of open αIIbβ3 integrin (PDB: 3FCU).

**Figure 4:**
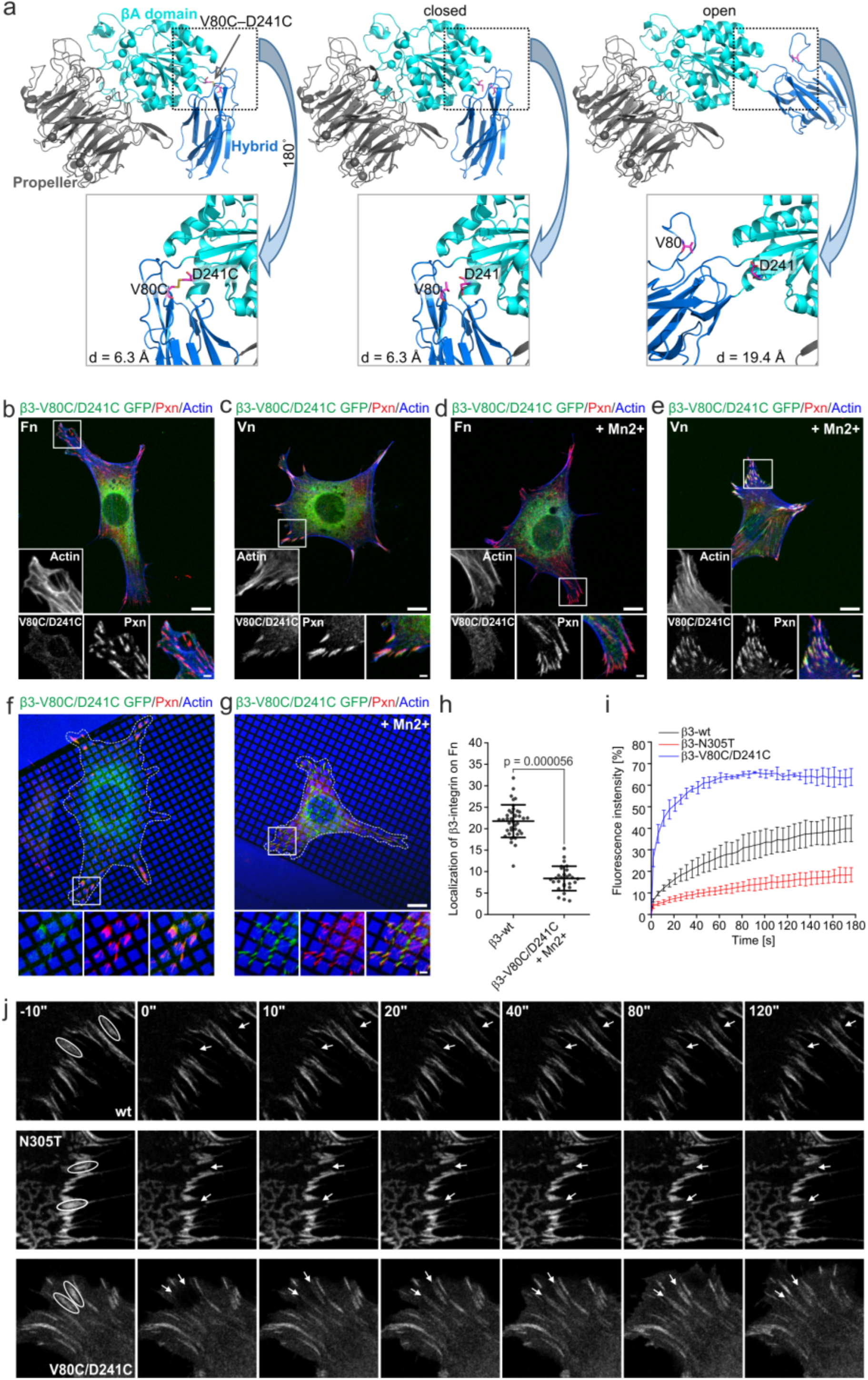
Extended-closed locked β3-V80C/D241C binds Vn but not Fn. (**a**) Structural analysis of the distance between V80 and D241 for extended-closed conformation of αVβ3 integrin (left; PBD: 4MMX) after introducing a V80C/D241C disulphide bridge, (middle) for the wt structure, or (right; PBD: 4MMX/3FCU) for the extended-open conformation. (**b-e**) NIH 3T3 cells transfected with β3-V80C/D241C GFP (green) cultured on the indicated ECM proteins for 2 h. 1 mM Mn^2+^ was added for the last 30 min where indicated. Cells were stained for paxillin (red) and actin (blue) after fixation. Please note the absence of αVβ3 integrin clustering on Fn and the increased localization of β3-V80C/D241C in adhesions on Vn for Mn^2^+ treated compared to untreated cells. (**f, g**) Cells were prepared as described in (**b-e**) except that they were cultured on Fn/Vn substrates. (**h**) Quantifications of colocalization of β3 GFP with Fn for cells treated as described g (β3-wt: replot of the data from Fig. 3g; β3-V80C/D241C + Mn2+: n = 29, N = 3). (**i, j**) NIH3T3 cells were transfected with the indicated plasmids and cultured on serum coated cover slips for 15 - 20 h. (**i**) FRAP measurement of αVβ3 integrin dynamics for the indicated conditions with (**j**) representative examples (β3-wt: n = 40, N = 3; β3-N305T: n = 48, N = 3; β3-V80C/D241C: n = 36, N = 3). (**b-g**) Scale bars: overview images 10 μm, zoom-ins 2 μm. White dashed lines indicate the cell outline. Fluorescent images were taken (**b-g**) with diffraction limited microscopy or (**j**) with confocal microscopy.

Next, we expressed this β3-V80C/D241C GFP in NIH 3T3 cells and cultured them on homogenous Fn or Vn (Fig. 4b, c). The GFP signal revealed clustering of β3-V80C/D241C into cell adhesions on Vn but no clustering at all on Fn. Opening disulfide bridges with dithiothreitol (DTT) allowed clustering into adhesions similar to β3-wt (Supplementary Fig. 4a, b, g). Additionally, incubating cells with Mn^2^+ failed to induce clustering of β3-V80C/D241C on Fn while it increased clustering on Vn (Fig. 4d, e). On Fn/Vn substrates, β3-V80C/D241C GFP showed also very dim adhesions that we could not quantify reliably due to high background fluoresence in the plasma membrane (Fig. 4f). However, a restriction of β3-V80C/D241C GFP localization on Vn was obvious. Incubating β3-V80C/D241C GFP expressing cells with Mn^2^+ increased the clustering of the mutated αVβ3 integrin while preserving the restriction to Vn (Fig. 4g, h). The increased fluorescent signal in focal adhesions compared to the plasma membrane in this experiment allowed a reliable analysis of β3-V80C/D241C GFP localization. The quantification revealed that the β3-V80C/D241C mutation has clear defects in Fn-compared to Vn-binding while our structural analysis confirmed that this mutation locks αVβ3 integrin in the extended-closed conformation. Additionally, adding the integrin inactivator Ca2+ to β3-wt GFP expressing cells reduced Fn-binding of αVβ3 integrin (Supplementary Fig. 3c, d) to a similar extent as contractility inhibition or the β3-V80C/D241C mutant. Therefore, we propose that αVβ3 integrin binds Vn initially in the extended-closed conformation, whereas αVβ3 integrin binds Fn only in the extended-open conformation.

We also compared the fluorescence recovery after photobleaching (FRAP) of β3-wt, β3-N305T, and β3-V80C/D241C, in order to understand the influence of these mutations on αVβ3 integrin turnover in adhesions (Fig. 4i, j). Interestingly, β3-V80C/D241C GFP showed very fast turnover, while β3-N305T GFP showed slow integrin exchange compared to β3-wt GFP as reported previously (Cluzel et al., 2005). Combined with our experiments on Fn/Vn substrates, this revealed that Fn binding of αVβ3 integrin correlates with reduced integrin turnover on Vn substrates. Control experiments with overexpression of FAK wt vs. FAK Y397F confirmed this correlation: Despite proper focal adhesion recruitment, FAK Y397F fails to recruit Src kinases causing increased focal adhesion size and reduced adhesion dynamics compared to FAK wt (Swaminathan et al., 2016, Hamadi et al., 2005, Schaller et al., 1994). At the same time, cells cultured on Fn/Vn substrates revealed increased colocalization of αVβ3 integrin with Fn when overexpressing FAK Y397F compared to FAK wt (Supplementary Fig. 4e, f, i). Thus, regulation of the exchange dynamics of αVβ3 integrin and αVβ3 integrin-mediated adhesions, emerged as an additional option to regulate Fn-binding of αVβ3 integrin.

#### MD simulation supported conformation-dependent Fn binding

The experiments with β3-N305T emphasized the influence of the extended-open conformation for Fn binding of αVβ3 integrin. To better understand the influence of different integrin conformations on Fn and Vn binding at a single protein level, we employed MD simulations. Because no suitable structural data for Vn was available, we used a CRGDC peptide as a proxy for a high-affinity ligand of αVβ3 integrin. We investigated the binding of the RGD-containing 10th domain of Fn (FnIII10; Supplementary Fig. 5a) and of the CRGDC peptide (Supplementary Fig. 5c) to extended conformations of αVβ3 integrin exhibiting either the open or closed headpiece. Interaction energies between αVβ3 integrin and the CRGDC peptide were stable and similar over the simulation time for the extended-closed integrin (Supplementary Fig. 5f); the extended-open conformation showed similar but less stable interaction energies (Supplementary Fig. 5g). In contrast, interaction energies for FnIII10 showed stable interaction with αVβ3 integrin only for the extended-open conformation (Supplementary Fig. 5b). Thus, these results support the model where FnIII10 - αVβ3 integrin binding is less stable in the extended-closed conformation, and becomes more stable once the integrin is in the extended-open form upon full hybrid domain swing-out. Binding of CRGDC, in contrast, appeared stable already for the extended-closed conformation of αVβ3 integrin. Additionally, structural analysis indicated that full-length Fn, compared to FnIII10, might show even higher preference for the extended-open conformation due to lower sterical hindrance compared to the extended-closed conformation (Supplementary Fig. 5d, e).

**Figure 5:**
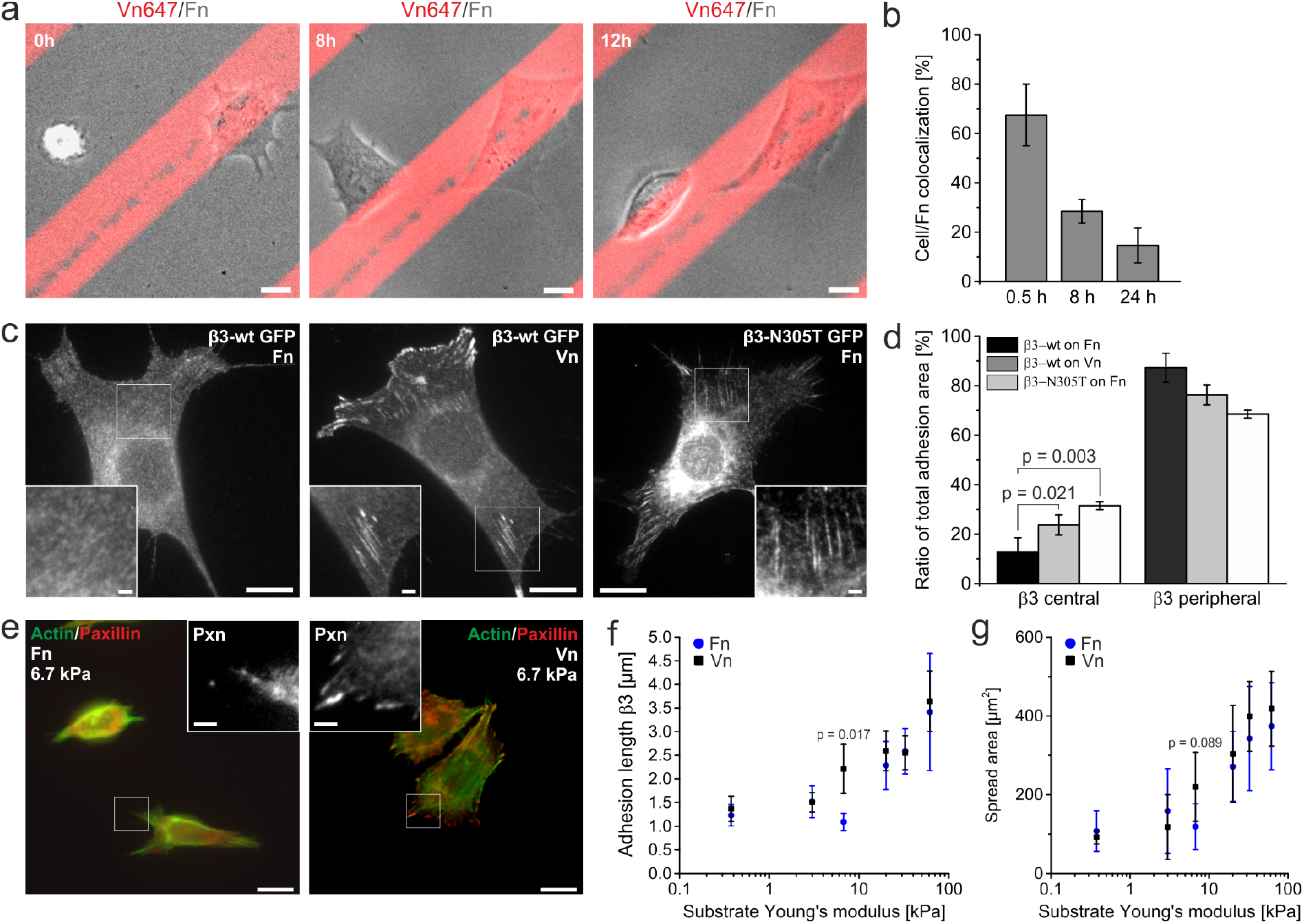
The preference of αVβ3 integrin for Vn influences cellular behavior. (**a**) GD25 cells were seeded onto stripe assays of stamped Vn (red; 10 μg/ml; 20 μm width) backfilled with Fn (10 μg/ml; 40 μm width). Alexa Fluor 647 labeled Vn was added to visualize Vn stripes. GD25 cells were visualized with phase contrast microscopy for 12 h. (**b**) Quantification of the colocalization of GD25 cells with Fn on Vn/Fn stripe assays at the indicated time points. The first time point was quantified based on phase contrast movies as described in (**a**) while 8 h and 24 h time points were calculated from experiments with cells cultured in the incubator, fixed, and stained for actin (0.5h: n = 123, N = 3; 8h: n = 17, N = 3; 24h: n = 15, N = 3) (**c**) NIH 3T3 cells were transfected with the indicated plasmid and cultured for 6 h on cover slips coated with the indicated ECM protein. (**d**) Quantification of the ratio of αVβ3 integrin-mediated adhesion area within a peripheral 2.5 μm frame around the cell contour compared to adhesions in the cell center (β3-wt on Fn: n = 23, N = 4; β3-wt on Vn: n = 26, N = 4; β3-N305T on Fn: n = 17, N = 3). (**e-g**) GD25wt were cultured on elastic polyacrylamide gels of the indicated Young’s modulus (E) and stained for paxillin (red) and actin (green). Gels were coated homogeneously with Vn or with Fn. (**e**) Cells on 6.7 kPa hydrogels showed less cell spreading and adhesion maturation on Fn coated substrates compared to Vn. (**f**) Length of paxillin-stained adhesions (longest 10% only) or (**g**) cell area was plotted against the Young’s modulus for cells on Vn (black data points) or Fn (blue data points; see Supplementary Table 1 for number of analyzed cells; p-values except the indicated: p > 0.1; see also Supplementary Table 2). All cells were imaged with diffraction limited microscopy. (**a, c, e**) Scale bar: 10 μm in overviews, 2 μm in zoom-ins.

#### Preference for Vn influences cell migration and mechanotransduction

Our results so far revealed a mechanism enabling αVβ3 integrin to differentiate between Fn and Vn based on the degree of the force-dependent hybrid domain swing-out. However, stable binding to both ligands will most likely result in the fully active extended-open conformation of αVβ3 integrin irrespective of the actual ligand present. Therefore, it is possible that the preference of αVβ3 integrin for Vn compared to Fn is compensated on a cellular level when only one ligand is present. Thus, we performed additional experiments to test this hypothesis. First, we analyzed cell migration of GD25 cells (no β1 integrin expression and therefore relying on αVβ3 integrin only) on cover slips coated with 10 μg/ml Fn or Vn with live cell imaging. Cell tracking revealed that cells on Fn migrated almost two times faster (vFn = 12.0 ± 3.08 μm/h) compared to cells migrating on Vn (vVn = 6.7 ± 0.39 μm/h; Supplementary Video 7). To understand how cell migration is influenced when GD25 cells can choose between Vn and Fn, we produced a Vn/Fn stripe assay (Vn: 20 μm width; Fn: 40 μm width). Live cell imaging over 12 h on these Fn/Vn stripe patterns revealed a consistent movement of cells from Fn towards Vn (Fig. 5a and Supplementary Video 8). To quantify this behavior, we measured the area of single cells covering Fn coated surfaces for cell cultures after different time points (Fig. 5b). Half an hour after seeding, cells covered Fn and Vn coated surfaces according to the geometrical coverage of the pattern indicating random distribution (1/3 Vn, 2/3 Fn, cell/Fn colocalization: 67.5%). With increasing time, more cells adhered to Vn (cell/Fn colocalization after 8 h: 28.4%; 24 h:14.6%).

Next, we investigated cellular behavior that have been described to be regulated by force-dependent mechanisms. First, we analyzed the subcellular localization of αVβ3 integrin. For cells cultured on Fn, it is well known that αVβ3 integrin dominates in peripheral, high-contractile focal adhesions whereas α5β1 integrin establishes central, low-contractile fibrillar adhesions (Zamir et al., 2000). Based on our data, it is straightforward to propose that this integrin distribution is caused by an interplay of extracellular ligands and mechanical load on the integrin: αVβ3 integrin fails to bind Fn in the cell center due to the low force levels in fibrillar adhesions. On Vn, in contrast, we expected that low mechanical load on αVβ3 integrin in the cell center is still compatible with Vn binding. We tested this by culturing β3-wt GFP transfected NIH 3T3 cells on homogenous Fn or Vn substrates for 6 h and analyzed the subcellular distribution of αVβ3 integrin. We frequently observed central αVβ3 integrin adhesions for cells cultured on Vn but rarely on Fn (Fig. 5c, d). In addition, we analyzed the β3-N305T mutation in this assay. The almost complete extended-open conformation of this mutation might allow αVβ3 integrin to bind Fn in the cell center despite the reduced mechanical load compared to the cell periphery. Indeed, β3-N305T GFP expressed in cells cultured on Fn now localized in central adhesions (Fig. 5c, d).

In a second set of experiments, we asked whether mechanotransduction of a cell is affected by the force-dependent ligand binding of αVβ3 integrin. We used again GD25 cells, seeded them on Fn or Vn coated hydrogels of different stiffnesses, and analyzed cell area and adhesion maturation as indicators of cellular mechanotransduction. Both ligands caused a similar sigmoidal increase of cell area and adhesion length with increasing gel stiffness (Fig. 5f, g). Cells on Vn showed adhesion maturation and increased cell spreading already at 6.7 kPa. Cells on Fn reached similar plateau values for both parameters, however, at substrates stiffer than 6.7 kPa (Fig. 5e).

To summarize, we observed that cellular behavior is regulated by the extracellular ligand of αVβ3 integrin. Cell migration experiments indicated that ligand preferences of αVβ3 integrin guide cells towards the high-affinity ligand Vn. More importantly, mechanotransduction is differently affected on Vn and Fn implying that force-dependent ligand-binding by αVβ3 integrin is not compensated on a cellular level.

#### Ligands of αVβ3 integrin belong to two different affinity regimes

Finally, we asked whether our findings for the interaction of αVβ3 integrin with Fn and Vn are also valid for other ligand combinations. We performed additional affinity measurements of αVβ3 integrin and its ligand fibrinogen (Fbg) revealing slightly higher affinity compared to Fn (Fbg: K_D_ = 8.15 ± 4.82 nM; Supplementary Fig. 3i, k). In addition, we produced binary choice substrates (Fig. 6g) to challenge αVβ3 integrin with Vn and Fbg (Fig. 6a), with Vn and osteopontin (Opn; Fig. 6b), or with Vn and thrombospondin (Tsp, Fig. 6c). Interestingly, αVβ3 integrin colocalized with Fbg (Fig. 6d) to a similar amount as it did with Fn on Fn/Vn substrates (colocalization to Fn: 16.5% on Fn/Vn, colocalization to Fbg 20.2% on Vn/Fbg). Opn caused a very different distribution with almost equal localization of αVβ3 integrin on both ligands (Fig. 6e, colocalization to Opn 50.4% on Vn/Opn). On Vn/Tsp, in contrast, αVβ3 integrin localized almost only on Vn (Fig. 6f, colocalization to Tsp 6.7% on Vn/Tsp) indicating that Tsp is no proper ligand for αVβ3 integrin in this context. Thus, Vn and Opn may present a class of high-affinity ligands for αVβ3 integrin compared to the low-affinity ligands Fn and Fbg.

**Figure 6:**
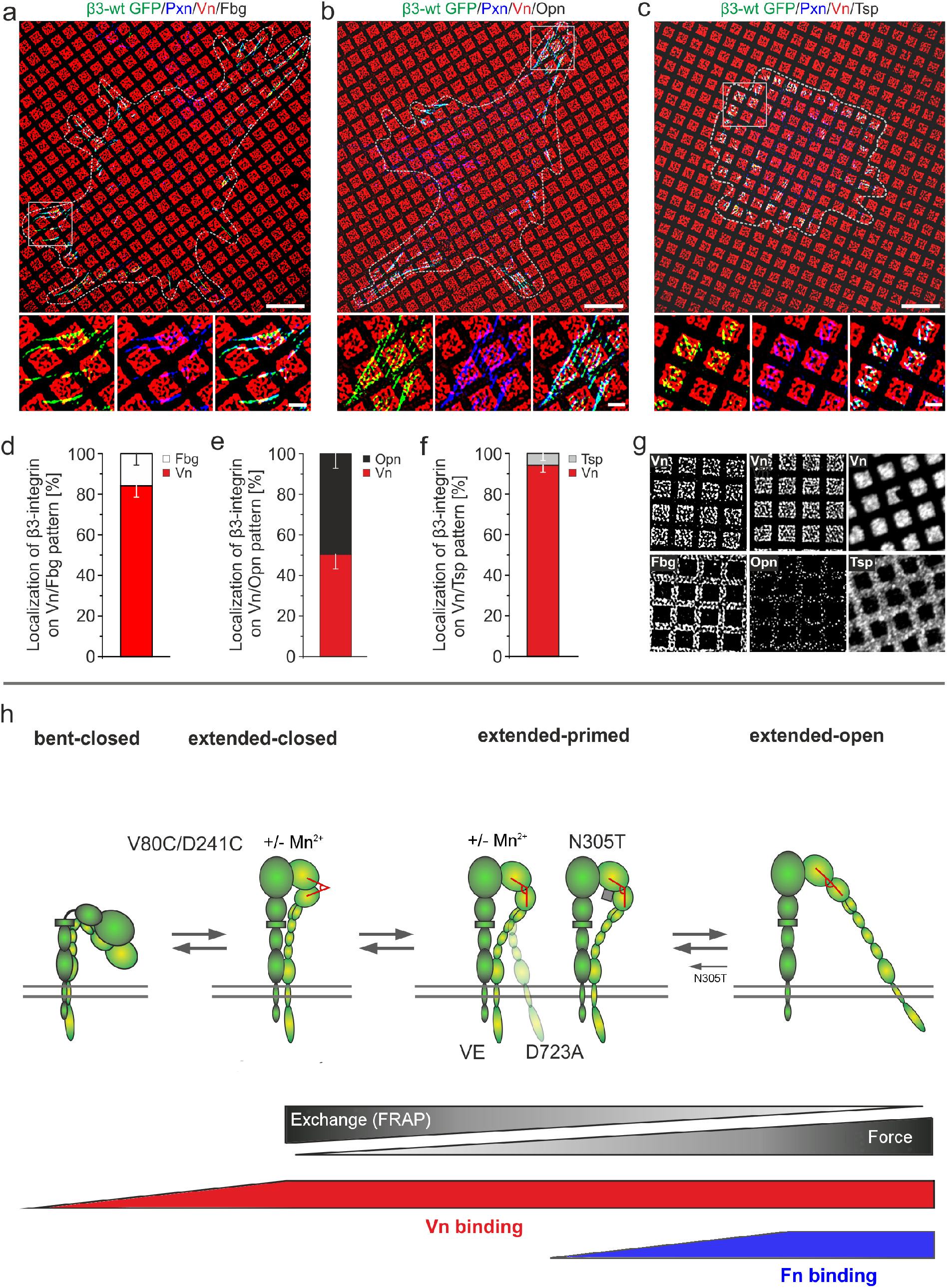
Vn and osteopontin (Opn), in contrast to Fn and fibrinogen (Fbg), are high-affinity ligands for αVβ3 integrin. (**a**) Vn squares (red) were stamped onto cover slips and the remaining area was backfilled with Fbg (black). NIH 3T3 cells transfected with β3-wt GFP integrin (green) cultured on Vn/Fbg substrates and immunostained for paxillin (blue). (**b**) Cell treated as in (**a**) but on a Vn/Opn or (**c**) a Vn/thrombospondin (Tsp) pattern. (**d-f**) Quantification of the colocalization of β3-wt GFP with indicated ECM proteins for cells cultured on Vn/Fbg (n = 57, N = 3), Vn/Opn (n = 45, N = 3), or Vn/Tsp (n = 34, N = 3). (**g**) Staining of Vn and the respective ECM proteins indicate successful protein separation for all binary choice substrates. Scale bars: (**a-c**) 10 μm in the overview, 2 μm in the zoom in. White dashed lines indicate cell outline. All images acquired with SR-SIM except Vn/Tsp pattern in g. (**h**) Model for force-dependent differential ligand binding of αVβ3: αVβ3 integrin is in an equilibrium between the bent and extended conformations. Extended αVβ3 integrins can be subdivided in a spectrum of several steps (Zhu and Springer, 2013). Integrin mutations can represent certain steps as for example β3-N305T without mechanical load resembles step 7 of 8. More generally, these intermediate extended conformations could be labeled extended-primed states in contrast to extended-closed and -opened conformations. FRAP measurements for several mutations indicate that increasing αVβ3 integrin activation correlates with decreasing exchange dynamics (Fig. 4i, (Cluzel et al., 2005)). On the other side, maximal αVβ3 activation requires mechanical load on the integrin that is, at the same time, necessary for Fn-but not Vn-binding. Mn^2^+ increases the affinity of the integrin head domain but fails to increase Fn-binding without additional conformational changes. Thus, mechanical forces favor the full αVβ3 integrin activation that enables binding to additional ligands and enhances thereby ligand promiscuity of αVβ3 integrin.

Additionally, we asked whether the closely related αVβ5 integrin also shows a force-dependent binding to Fn. We transfected NIH 3T3 cells with β5-wt GFP and cultured these cells on Fn/Vn substrates. Comparing β5-wt GFP with anti-paxillin staining revealed two classes of adhesions. The majority of αVβ5 integrin clustered in the cell center without recruiting paxillin (Supplementary Fig. 4c). Only a subset of αVβ5 integrin colocalized with paxillin in the cell periphery. We quantified the colocalization of these peripheral αVβ5 integrin adhesions with Fn for control conditions and blebbistatin treated cells (Supplementary Fig. 4c, d, h). These measurements also showed a force-dependent binding of αVβ5 integrin to Fn (αVβ5 integrin + DMSO: (36.5 ± 13.47)%; αVβ5 integrin + 5 μM blebbistatin: (16.1 ± 4.84)%). Based on these findings it appears that force-dependent ligand binding is a general mechanism regulating the interaction of integrins with ECM-ligands.

### Discussion

Here we have analyzed the interaction of αVβ3 integrin with different ligands on sub-cellular binary choice substrates. We found that αVβ3 integrin prefers binding to Vn under a wide range of conditions. Binding to Fn, in contrast, required mechanical load on the integrin. Combining experiments with integrin mutations and MD simulations revealed that this differential ligand binding is coupled to different integrin conformations (Fig. 6h). The extended-closed conformation of αVβ3 integrin binds to Vn but not efficiently to Fn. Only the extended-open conformation due to a force-mediated complete hybrid domain swing-out enabled αVβ3 integrin to bind Fn. Thus, force-mediated conformational changes regulate the ligand promiscuity of αVβ3 integrin. We demonstrated that these findings have consequences for cellular behavior during migration and mechanotransduction.

### Fn vs. Vn preference of αVβ3 integrin

The preference of αVβ3 integrin for Vn compared to Fn is apparent in all our experiments. This might appear trivial given the measured *in vitro* affinities of αVβ3 integrin for Vn compared to Fn. However, this explanation fails to explain *in cellula* results like the exclusive initiation of αVβ3 integrin-mediated adhesions on Vn while αVβ3 integrin binds Fn in maturing focal adhesions. The reduced Fn-binding after contractility inhibition also points to a more complex regulation of αVβ3 integrin ligand-binding. We interpret these results as a consequence of force-dependent increase in Fn binding of αVβ3 integrin. We had no access to a single-protein method that would have allowed us to measure force-dependent affinity changes for single αVβ3 integrins. Instead, our experimental results are based on ensemble measurements revealing equilibrium changes of a population of integrins. Accordingly, a lack of adhesion initiation by αVβ3 integrin on Fn is still compatible with transient binding of αVβ3 integrins to Fn on spatial and temporal scales below the resolution of the methods we used. Thus, our interpretation of ligand-αVβ3 integrin interactions are based on the binding of many integrins over seconds to their ligands and not on affinity measurements of single integrins and ligands. Yet, we think that a force-dependent change in affinity of single αVβ3 integrins is the most plausible explanation for our results. Avidity instead of affinity changes or differential recruitment of adapter proteins could be possible alternative explanations. In fact, we showed that vinculin recruitment is needed for increased Fn-binding of αVβ3 integrin. However, to date vinculin is best characterized as a transmitter of force from actin to the talin-integrin axis (Humphries et al., 2007, Rahikainen et al., 2017, Elosegui-Artola et al., 2016). Therefore, we conclude that the effect of vinculin in our experiments is best explained by its role as a force-transmitter. Additionally, recruitment of adhesome proteins to focal adhesions seemed rather unaffected by a vinculin knockout (Thievessen et al., 2013). Avidity regulation on the other hand, is also not easily compatible with our data. Nascent adhesions are the smallest clusters of αVβ3 integrin we could resolve. Single localization techniques estimated 40-50 integrins in such clusters (Shroff et al., 2008, Changede et al., 2015). Therefore, the differential ligand binding of αVβ3 integrin that we observed during adhesion maturation occurred on a range of 40 up to probably hundreds of integrins in focal adhesions. In contrast, affinity vs. avidity regulation is rather discussed for a range of one vs. ten integrins (Abrams et al., 1994). These pitfalls of alternative explanations let us favor a hypothesis of force-regulated differential affinity change of αVβ3 integrin (Fig. 6h). More importantly, our data obtained with different αVβ3 integrin mutations, structural analysis, and comparisons with MD simulations offer a mechanistic explanation supporting this hypothesis.

### The active αVβ3 integrin for binding Vn is not the active integrin for binding Fn

Live cell imaging and contractility inhibition indicated that retrograde actin flow and cellular contractility might influence the ligand binding of αVβ3 integrin. In fact, MD simulations predicted that mechanical load on an integrin, imitating retrograde actin flow, should favor the hybrid domain swing-out and therefore the extended-open conformation (Zhu et al., 2008, Puklin-Faucher et al., 2006). Zhu and colleagues concluded that mechanical forces activate integrins (Zhu et al., 2008). Integrin activation can be defined on a structural level and on a functional level. The latter definition is straightforward with ligand binding indicating an active integrin while the inactive integrin fails to bind a ligand. For the structural definition, it is probably not controversial to define a bent integrin as the conformation of the inactive and the extended conformation as the active integrin (Campbell and Humphries, 2011). These structures are also described as low-affinity (bent integrin) and high-affinity (extended integrin) conformations indicating the probability of ligand binding (Kong et al., 2009, Zhu et al., 2008). In this way, the link between structural and functional definition can be established.

However, some integrins show extended conformations that can be subdivided into extended-closed and extended-open. This is, for example, the case for αVβ3 integrin. Whether this is also the case for α5β1 integrin is still debated (Su et al., 2016, Miyazaki et al., 2018). These subclasses of αVβ3 integrin make it less clear whether extended-closed or extended-open αVβ3 integrin should be considered as the active conformation. Our data offered a surprising answer to this question by showing that the active αVβ3 integrin (functional definition) has different conformations for different ligands. Fn binding only to the extended-open αVβ3 integrin defines this conformation as the active integrin structure. In contrast, Vn binds already to the extended-closed conformation rendering the inactive conformation for Fn-binding to the active conformation for Vn-binding. This rather complex and situation-dependent structure-function relationship of αVβ3 integrin might also be relevant for the development of high-affinity inhibitors that should bind without activating the integrin. For such a purpose, it will also be necessary to better understand the molecular detail how ligands bind, or do not bind, to the binding pocket for different conformations of αVβ3 integrin. Our data indicate that the influence of mechanical forces at the binding pocket can alter the affinity in a ligand specific manner. An interesting aspect of this issue was recently postulated by Cormier and colleagues who argued that integrin activation could also be about improved accessibility of the RGD peptide of a ligand to the binding pocket in the αVβ3 integrin headpiece (Cormier et al., 2018). Interestingly, Fn presents its RGD peptide in a rather short loop implying that accessibility to the αVβ3 integrin binding pocket could indeed be a limiting step for this ligand.

Such a scenario would also explain the limited effects of Mn^2^+ treatment on Fn-binding by αVβ3 integrin assuming that Mn^2^+ increases the affinity for the RGD peptide but does not change the accessibility of Fn to the αVβ3 integrin binding pocket (Fig. 6h). Also, it was reported that Mn^2^+ in absence of a ligand failed to cause a conformational shift to active conformations for the closely related αIIbβ3 integrin (Zhu et al., 2013).

### The vitronectin receptor and the cell

Cell migration relies on stable adhesion to the substrate involving an extension phase and generation of new adhesions underneath lamellipodia or filopodia while subsequent actomyosin forces pull on these anchors to push the cell body forward compared to the substrate (Giannone et al., 2007). Therefore, adhesions have to switch from a short lived exploratory mode (nascent adhesions) to a more stable sessile mode (focal adhesions) (Choi et al., 2008). αVβ3 integrin (Elosegui-Artola et al., 2016) and α5β1 integrin (Kong et al., 2009, Friedland et al., 2009) support this process by increasing the lifetime of their bond to Fn when force is applied (i.e. a catch-bond). However, adhesions also have to lose their contact to the substrate to allow the cell body to move forward. This can occur by weakening of adhesions during translocation to the cell center (Zamir et al., 2000, Sun et al., 2016) and by disassembly of adhesions (Wehrle-Haller, 2012). Our Fn/Vn stripe assay (Fig. 5a) and our analysis of subcellular localization of αVβ3 integrin revealed that the respective ligand of an integrin can be decisive for adhesion organization and cell migration. Faster off-rates of the αVβ3 integrin-Fn bond (Fig. 1h) allow a faster cycle of binding and unbinding compared to Vn. Additionally, αVβ3 integrin builds less adhesions in total (Supplementary Fig. 2e) and specifically lower numbers of central adhesions on Fn (Fig. 5d). This is accompanied by less adhesive forces on Fn compared to Vn (Fig. 1j). In total, this leads to a more exploratory, faster migrating cell on Fn (Supplementary Video 7, 8). On Vn, in contrast, αVβ3 integrin causes higher adhesive forces, more adhesion area per cell, and sticks to Vn even under low mechanical load explaining why cells moved from Fn away towards Vn. This raises the interesting option that cell migration towards Vn from the blood plasma contributes to wound healing by attracting αVβ3 integrin-expressing cells to a site of coagulated serum proteins. We believe it might also be relevant for physiological settings that the Fn-binding capacity of αVβ3 integrin is not only regulated by forces but also by adhesion dynamics (Fig. 6h). Therefore, an αVβ3 integrin expressing cell would not be at the mercy of the surrounding tissue stiffness for binding, or not binding, to Fn. Instead, adhesion dynamics, regulated by multiple signaling cascades, offer an opportunity to tune the binding of αVβ3 integrin to Fn and to integrate mechanical and biochemical signals. On the other hand, assuming that the bond between a soluble ligand and an integrin experiences low mechanical load, our results can also explain the inability of αVβ3 integrin to bind soluble Fn (Huveneers et al., 2008) avoiding the clogging of surface exposed αVβ3 integrin by Fn from the blood plasma. Experiments with cells cultured on hydrogels of different stiffnesses revealed a mechanotransduction on a sigmoidal basis, i.e. an on-off mechanoswitch, as reported by others (Elosegui-Artola et al., 2016). Interestingly, GD25 cells on Vn coated gels exhibited increased αVβ3 integrin-mediated adhesion length at an intermediate gel stiffness (E = 6.73 kPa) where cells cultured on Fn coated gels still failed to establish mature adhesions. We showed that αVβ3 integrin establishes bonds to Vn, but not to Fn, under low-contractility conditions. Thus, assuming that softer gels lead to lower mechanical load on αVβ3 integrin, our findings are able to explain the differential mechanotransduction of cells on two different ligands.

In total, we showed that the same integrin, αVβ3, can behave differently and cause different cellular phenotypes dependent on the ligand it binds to. On the other side, intra- and extracellular parameters can affect the ability of αVβ3 integrin to select between different ligands. Interestingly, αVβ5 integrin was recently shown to contribute to non-canonical integrin functions either in so called reticular adhesions or by associating with clathrin containing structures (Zuidema et al., 2018, Lock et al., 2018, Baschieri et al., 2018). The switch between these structures and focal adhesions is regulated by different intra- and extracellular parameters including actomyosin-contractility. Moreover, a recent report showed that α4β7 integrin selects between VCAM-1 and MAdCAM-1 dependent on the integrin conformation induced by different chemokines (Wang et al., 2018). This indicates that at least some integrins can change their ligand selectivity and/or their function based on their conformation regulated by environmental factors. This proposes an additional, emerging way to control integrin behavior.

## Supporting information

## Acknowledgements

We thank Marc Hippler for help with atomic force microscopy measurements, Melanie Merkel for help with the analysis of de novo clusters, Deepthy Kavungal for help with cell migration assays, and the Bioimaging core facility at CMU, University of Geneva. This work was supported by the German Research Foundation (Karlsruhe School of Optics and Photonics (KSOP) to M. Bachmann) and by the Baden-Württemberg Stiftung (State Scholarship to M. Schäfer). We thank Academy of Finland for financial support (grant no. 290506 to V. Hytönen) and acknowledge CSC – IT center for science for computational resources and EDUFI (former CIMO) for postdoctoral fellowship for V. Mykuliak. Swiss National Science Foundation supported the work of M. Ripamonti and B. Wehrle-Haller (31003A_166384). The authors declare no competing financial interests.

## Author contributions

M. Bachmann, B. Wehrle-Haller, and M. Bastmeyer conceived the study. M. Bachmann, M. Schäfer, V. Hytönen, B. Wehrle-Haller, and M. Bastmeyer designed the experiments, M. Bachmann, M. Schäfer, V. Mykuliak, M. Ripamonti, L. Heiser, K. Weißenbruch, S. Krübel, and C. M. Franz performed and analyzed the experiments. M. Bachmann, M. Schäfer, B. Wehrle-Haller, and M. Bastmeyer wrote the paper. All authors discussed the results and implications and commented the manuscript drafts.

---

**Supplementary Figure 1:**
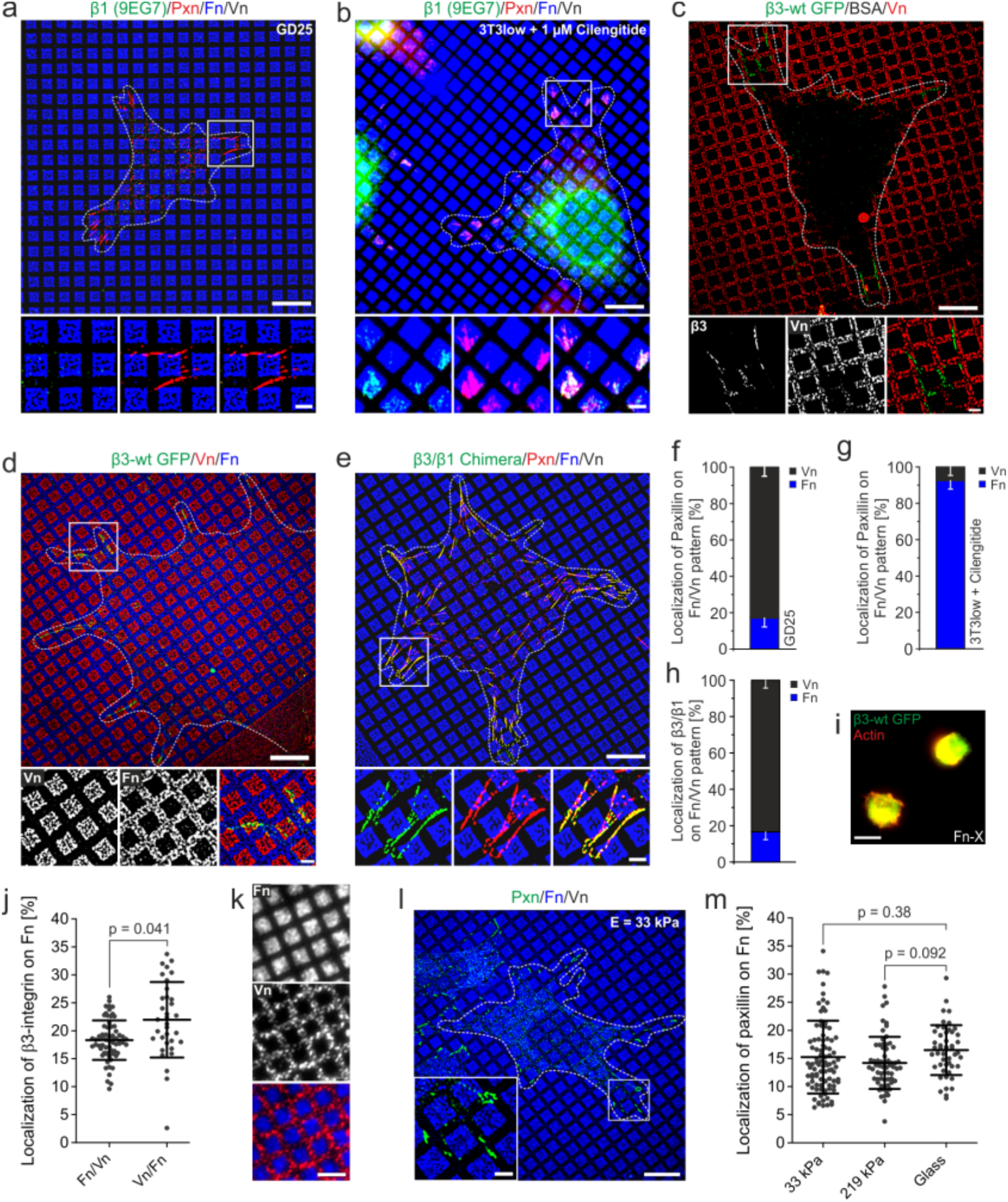
(**a**) GD25 cells cultured on Fn/Vn substrates (Fn in blue) and immunostained for β1 integrin (green) and paxillin (red). GD25 cells express no β1 integrin (shown by the immunostaining for β1 integrin; left zoom in and merge with paxillin on the right). (**b**) NIH 3T3 cells cultured on Fn/Vn substrates (Fn in blue) and immunostained for β1 integrin (green) and paxillin (Pxn; red). Cilengitide, an αVβ3 integrin inhibitor, was added 30 min before fixation. Paxillin is restricted to Fn. (**c**) Bovine serum albumin (BSA) was stamped and backfilled with Vn (red; labeled by immunostaining). NIH 3T3 cells transfected with β3-wt GFP integrin were cultured on these substrates. Zoom-ins show that clustered αVβ3 integrin is not present on BSA. (**d**) Vn labeled with Alexa Fluor 568 (red) was stamped and then overlaid with Alexa Fluor 647 labeled Fn (blue). NIH 3T3 cells were transfected with β3-wt integrin (green) and cultured on these Vn/Fn substrates. (**e**) NIH 3T3 cells were transfected with a β3/β1 integrin chimera (green) with the extracellular part of β3 integrin and the transmembrane and cytoplasmic part of β1 integrin. These cells were cultured on Fn/Vn substrates (Fn in blue) and immunostained for paxillin (red). (**f**) Quantification of paxillin colocalization with Fn in GD25 cells treated as described in a (n = 36, N = 3). (**g**) Quantification of paxillin colocalization with Fn in NIH3T3 cells treated as described in b (n = 27, N = 4). (**h**) Quantification of β3/β1 integrin chimera colocalization with Fn in NIH3T3 cells treated as described in e (n = 38, N = 3). (**i**) NIH 3T3 cells transfected with β3-wt GFP integrin cultured on heat-inactivated Fn (Fn-X) that was used together with native Fn during stamping (see Material & Methods). Cells are unable to spread and to cluster integrins on this substrate. (**j**) Quantification of β3 integrin colocalization with Fn in NIH3T3 cells treated as described in d. Category “Fn/Vn” is a replot of Fig. 1d (Vn/Fn: n = 38, N = 3). Please note that, besides a significant difference, β3-wt GFP preferentially localizes on Vn coated areas irrespective of the stamping procedure. (**k**) Protein separation of Fn/Vn patterns tested after transfer of the pattern to polyacrylamide gels by immunostaining for Fn (blue) and Vn (red). (l) Immunostained GD25 cell (paxillin: green; Fn: blue) on a gel with a Young’s modulus of E = 33 kPa functionalized with Fn/Vn patterns. The zoom-in highlights the preferred localization of adhesions on Vn. (**m**) Quantification of the colocalization of paxillin with Fn for cells grown on Fn/Vn pattern printed on glass (n = 36, N = 3), gels with E = 33 kPa (n = 93, N = 3), or gels with E = 219 kPa (n = 64, N = 3). (**a-e, l**) White dashed lines in overview images indicate cell outlines. Scale bar: 10 μm in the overview and 2 μm for the zoom-in and in (**k**). All fluorescent images except (**b, i, and k**) were acquired with SR-SIM.

**Supplementary Figure 2:**
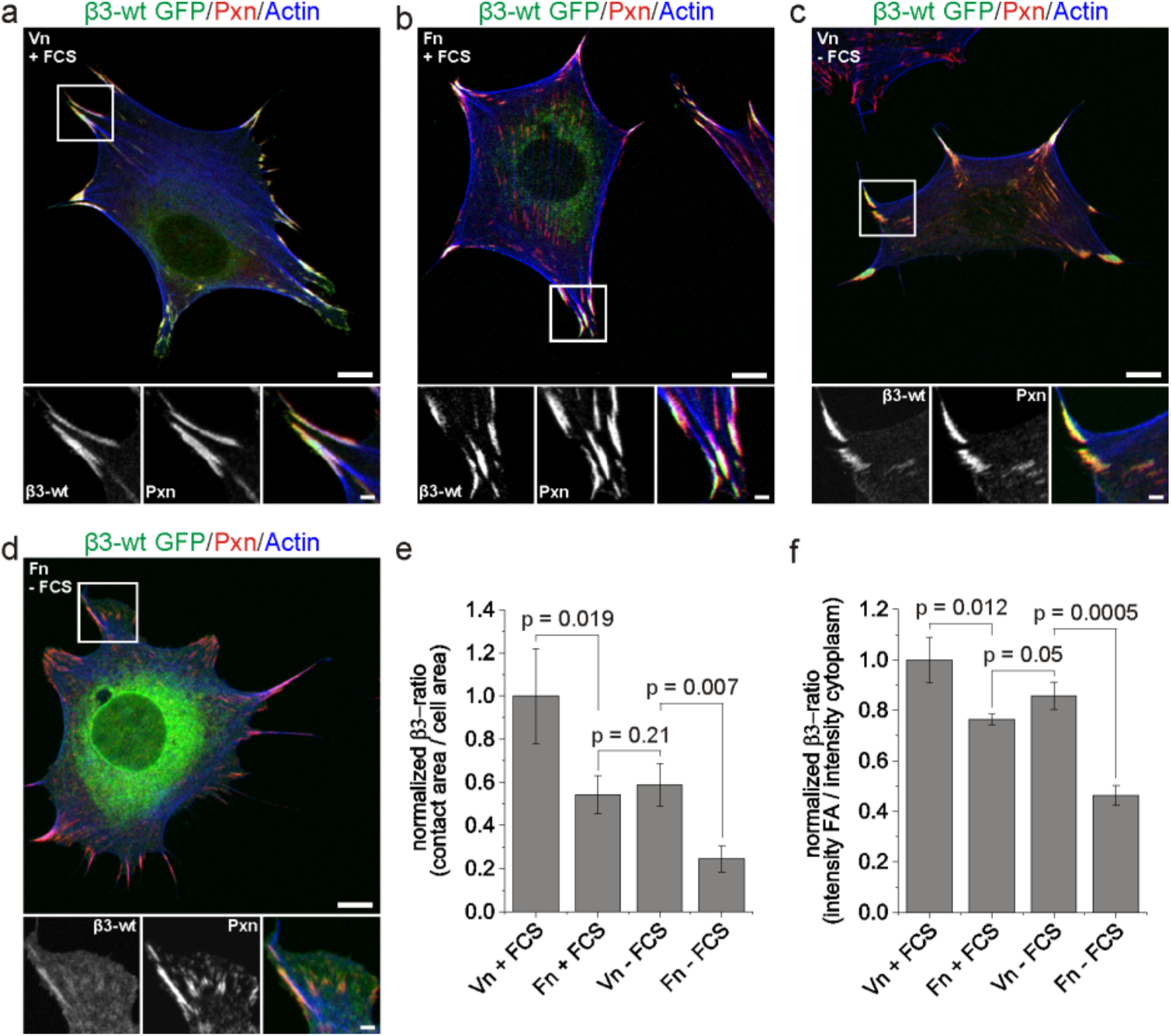
αVβ3 integrin favors Vn compared to Fn on homogenous substrates. (**a-d**) NIH 3T3 cells transfected with β3-wt GFP (green) cultured on homogenously coated substrates (Vn or Fn as indicated) and stained for anti-paxillin (Pxn; red) and actin (blue). Throughout the experiment, fetal calf serum (FCS) was present in the medium at 10% v/v (+FCS) or not (-FCS). (**e**) Quantification of the size ratio of β3-wt GFP mediated cell-matrix adhesions compared to the cell area for the conditions described in a-d. Ratios are normalized to the ratio for Vn + FCS (Vn+FCS: n = 67, Fn+FCS: n = 69, Vn-FCS: n = 70, Fn-FCS: n = 67; N = 3 in all cases). (**f**) Quantification of the intensity ratio of β3-wt GFP mediated cell-matrix adhesions compared to the plasma membrane around the adhesions for the conditions described in a-d. Ratios are normalized to the ratio for Vn + FCS (same cells as in e were used for quantification; N = 3 in all cases). All images were acquired with diffraction limited microscopy. Scale bars: 10 μm in the overview, 2 μm in the zoom-ins.

**Supplementary Figure 3:**
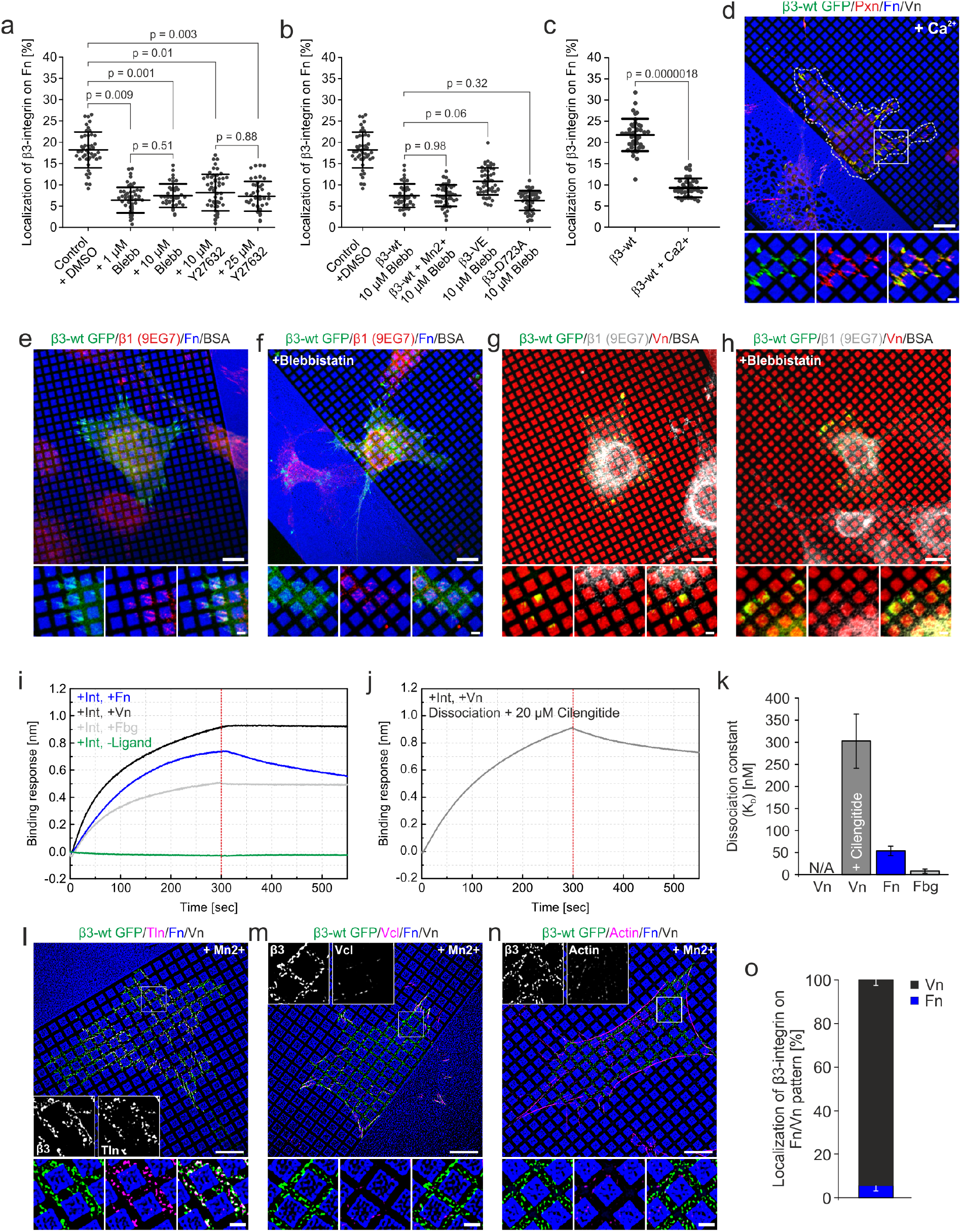
(**a**) Quantification of β3 integrin colocalization with Fn in NIH3T3 cells transfected with β3-wt integrin and cultured on Fn/Vn substrates. Cells cultured in presence of blebbistatin or Y27632 at the indicated concentrations. Values of 10 μM inhibitor concentration and DMSO control are replotted from Fig. 2g (+ 1 μM Blebb: n = 51, N = 3; + 25 μM Y27632: n = 56, N = 3). (**b**) Quantification of β3 integrin colocalization for the indicated mutations with Fn in NIH3T3 cells transfected with β3-wt integrin and cultured on Fn/Vn substrates. 30 min before fixation blebbistatin (and Mn^2^+ where indicated) was added at the indicated concentrations. Values of DMSO control and “β3-wt +10 μM Blebb” are a replot of Fig. 2g (β3-VE+10 μM Blebb: n = 55, N = 3; β3-wt+Mn^2^+ +10 μM Blebb: n = 45, N = 3; β3-D723A+10 μM Blebb: n = 49, N = 3). (**c**) Quantification of β3 integrin colocalization with Fn for cells treated as described in Fig. 1c (β3-wt) or in d (β3-wt +Ca2+). (**d**) NIH 3T3 cells transfected with β3-wt GFP (green) cultured on Fn/Vn substrates for 2 h. 1 mM Ca^2^+ was added for the last 30 min. Cells were stained for paxillin (red) and actin (blue) after fixation. (**e-h**) NIH 3T3 cells transfected with β3-wt integrin GFP (green) and cultured on (**e, f**) Fn(blue)/BSA or (**g, h**) Vn(red)/BSA patterns. 5 μM blebbistatin was present during the experiment where indicated. After fixation cells were immunostained for active β1 integrin by staining with clone 9EG7 antibodies (red in **e, f**; white in **g, h**). (**i**) Association (0 – 300 s) and dissociation (300 – 550 s) curves for the interaction of the indicated ECM proteins with purified αVβ3 integrin. The measurements were performed with biolayer interferometry (representative curves from n = 9, N = 3 measurements for each condition). (**j**) Association and dissociation curve for Vn interaction with purified αVβ3 integrin. Cilengitide was present in the dissociation buffer at 20 μM concentration (representative curve from n = 9, N = 3 measurements). (**k**) Quantification of dissociation constants for the indicated conditions based on the graphs shown in **i, j**. (**l-n**) NIH 3T3 cells transfected with β3-wt GFP (green), cultured on Fn/Vn substrates and treated with 1mM Mn^2^+ 30 min before fixation. Additionally, cells were immunostained for (**l**) talin, (**m**) vinculin, or (**n**) for actin (magenta). (**o**) Quantification of β3-wt GFP in central areas of Mn^2^+ treated cells cultured as described in **l-n** but immunostained for paxillin. Only β3-wt GFP not colocalizing with paxillin was used for analysis to ensure exclusive measurement of nascent integrin clusters (n = 42; N = 3). (**d-h, l-n**) Scale bars: 10 μm in the overview, 2 μm in the zoom-ins. (**d-h**) were acquired with diffraction-limited microscopy, (**l-n**) with SR-SIM.

**Supplementary Figure 4:**
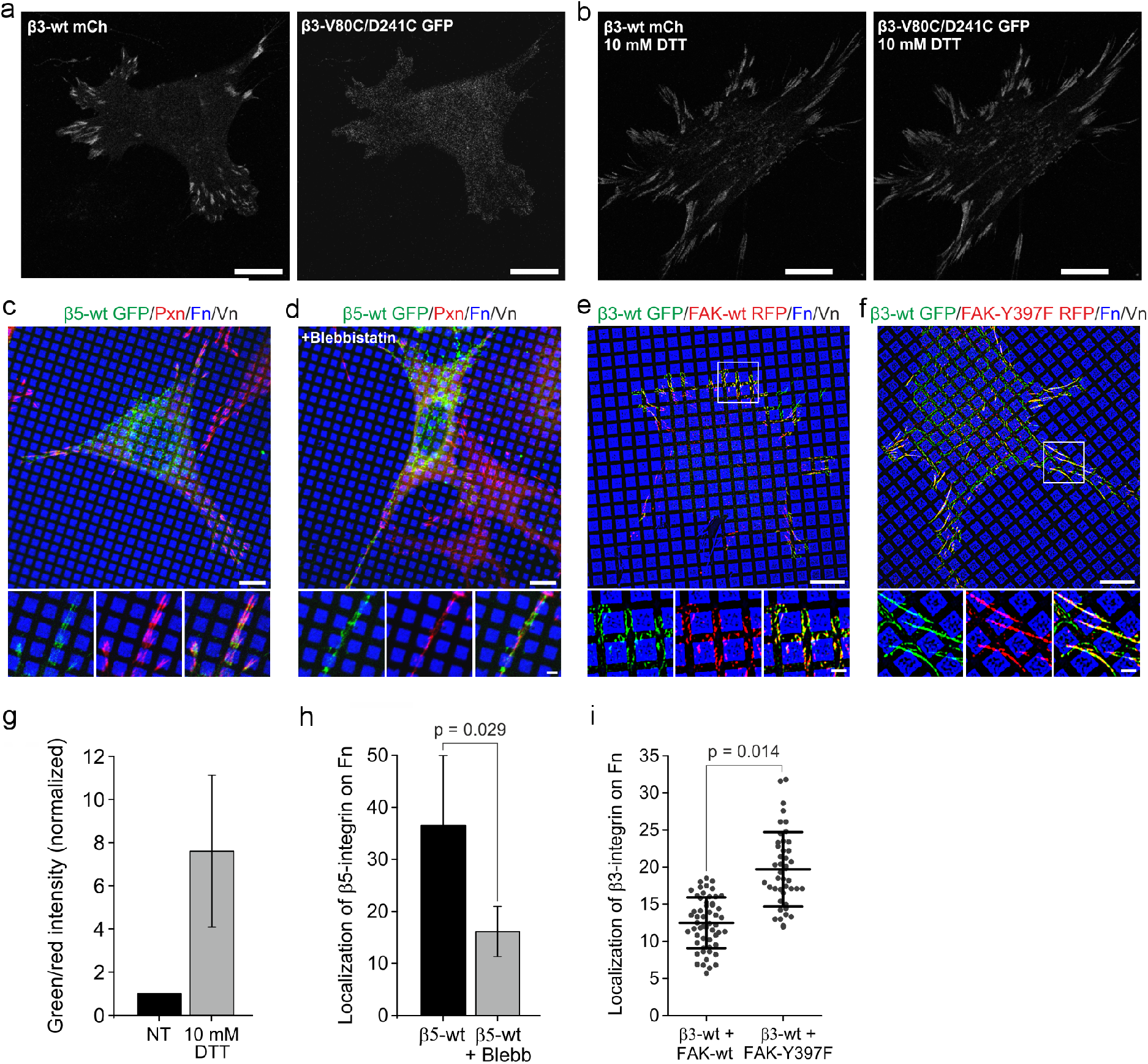
(**a, b**) NIH 3T3 cells transfected with β3-wt mCherry and β3-V80C/D241C GFP and cultured with 10% FCS on serum coated coverslips for 17 h. (**b**) 10 mM dithiothreitol (DTT) was added 10 min before fixation to open disulphide bridges. (**c, d**) NIH 3T3 cells transfected with β5-wt GFP (green) and cultured on Fn/Vn substrates as described but prepared with reduced Vn (0.1 μg/ml). Cells were fixed after 2 h and immunostained for paxillin (red). 1 μM blebbistatin was present when indicated. (**e, f**) NIH 3T3 cells were transfected with β3-wt GFP (green) and (**e**) FAK-wt RFP or (**f**) FAK-Y397F RFP (red) and cultured on Fn/Vn substrates. FAK signal was in both cases increased by anti-HA tag staining. (**g**) Quantification of fluorescence intensity of GFP/mCherry in focal adhesions for cells treated as described in **a, b**. DTT treatment increases the signal of β3-V80C/D241C compared to β3-wt. Intensity ratio was normalized to the non-treated (NT) control. (**h**) Quantification of β5-wt GFP colocalization with Fn for cells as described in **c, d** (β5-wt: n = 26, N = 5; β5-wt + Blebb: n = 24, N = 4). (**i**) Quantification of β3-wt GFP colocalization with Fn for cells as described in **e, f** (β3-wt + FAK-wt: n = 55, N = 3; β3-wt + FAK Y397F: n = 44, N = 3). Scale bar: 10 μm in the overview and 2 μm for the zoom-in. All fluorescent images were acquired with diffraction limited microscopy.

**Supplementary Figure 5:**
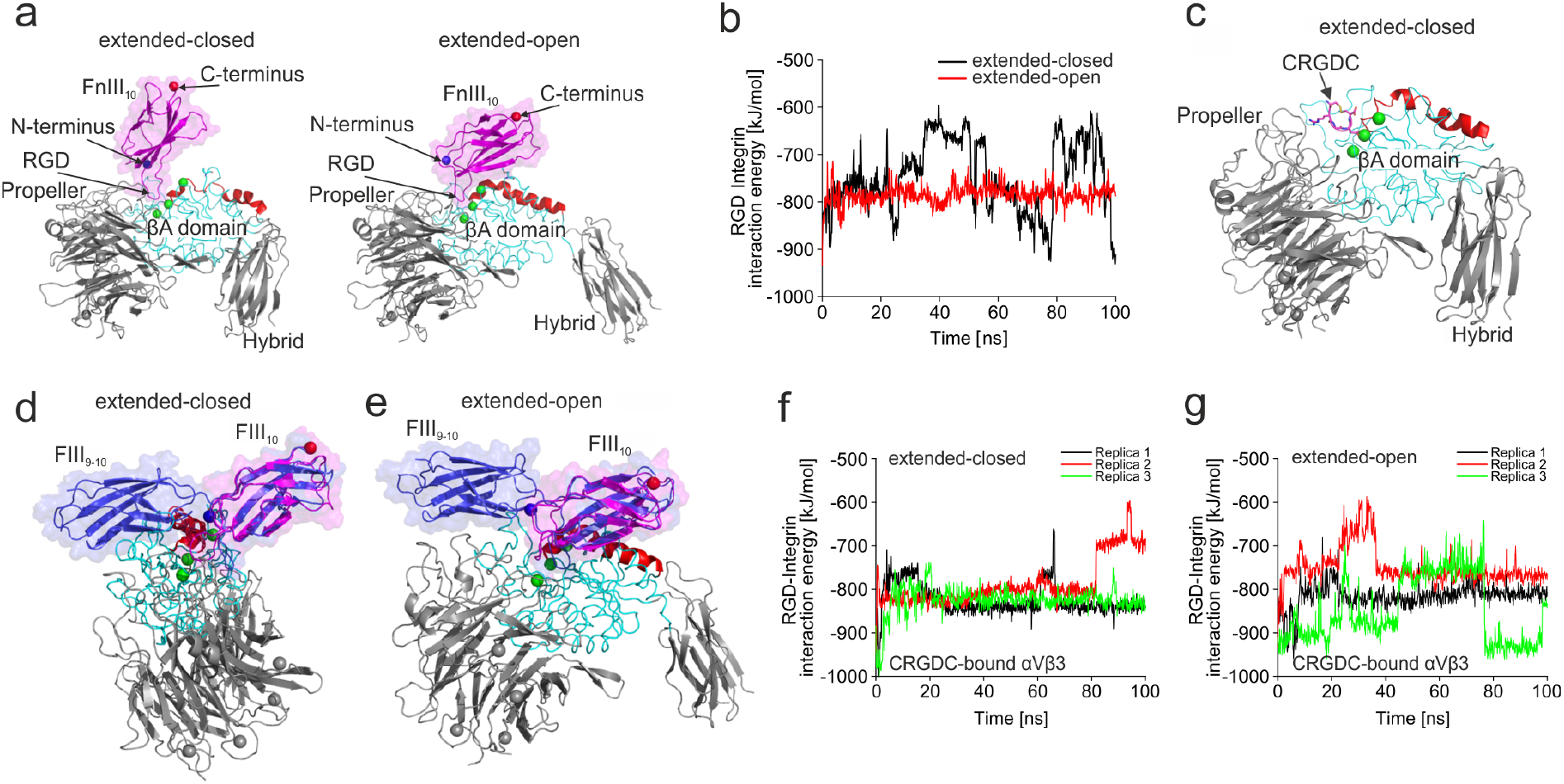
MD simulations support that Fn binds to extended-open αVβ3 integrin. (**a**) Snapshot of extended-closed (PBD: 4MMX) and extended-open (prepared as described in Fig. 4a; PBD: 4MMX/3FCU) conformation of FnIII10-bound αVβ3 integrin of a MD simulation. After initial 30 ns-simulations for FnIII10-binding, we selected the best replicas based on FnIII10-orientation and interaction energy between the RGD-sequence of FnIII10 and αVβ3 integrin, and proceeded for 100 ns simulations. Please note that the orientation of FnIII10-binding to the extended-closed conformation is different as compared to the one observed for FnIII10 bound to the extended-open conformation, indicating differences in the binding capacity of the respective integrin conformations. (**b**) Interaction energy between the RGD motif of FnIII10 and integrin over time of the MD simulation. (**c**) Snapshot of extended-closed conformation of CRGDC-bound αVβ3 integrin taken after 100 ns MD simulation. The FnIII10-bound integrin structures were used as templates to prepare the CRGDC-bound αVβ3 structures. (**d, e**) Overlay of extended-closed and extended-open conformation from Supplementary Fig. 5a with a Fn structure consisting of 9^th^ and 10^th^ domain of full-length Fn (FnIII9-10; blue). Please note steric clashes for the extended-closed conformation. Loops of the βA domain of β3 integrin (cyan) overlap with the 9^th^ domain of Fn. This is not the case for the extended-open conformation. (**f, g**) RGD-integrin interaction energies for CRGDC-bound αVβ3 integrin in extended-closed or extended-open conformation from three independent MD simulations.

## Supplementary Videos

Supplementary Video 1: NIH 3T3 cell expressing β3-wt GFP (green) was monitored during spreading on Fn/Vn substrate (Fn in blue) with live-cell SR-SIM with 1 frame per minute. The imaging medium contained 10% FCS. See also Fig. 1g.

Supplementary Video 2: Zoom-in of Supplementary Video 1.

Supplementary Video 3: NIH 3T3 cell expressing β3-N305T GFP (white) was monitored during spreading on Fn/Vn substrate (Fn in blue) with live-cell SR-SIM with 1 frame per minute. The imaging medium contained 10% FCS. See also Fig. 3j.

Supplementary Video 4: Zoom-in of Supplementary Video 3.

Supplementary Video 5: NIH 3T3 cell expressing β3-VE GFP (white) was monitored during spreading on Fn/Vn substrate (Fn in blue) with live-cell SR-SIM with 1 frame per minute. The imaging medium contained 10% FCS.

Supplementary Video 6: Zoom-in of Supplementary Video 5.

Supplementary Video 7: GD25 cells cultured on homogenous Fn (left) or Vn (right) (10 μg/ml) and imaged with phase contrast microscopy with 1 frame per 20 minutes. 34 cells from three independent experiments were analyzed in both cases. The imaging medium contained 1% FCS.

Supplementary Video 8: GD25 cells cultured on Vn/Fn stripes and imaged with phase contrast microscopy with 1 frame per 20 minutes. Stripes of Vn (red) have a width of 20 μm. The imaging medium contained no FCS.

**Supplementary Table 1:** Polyacrylamide gel rigidities measured with AFM. Number of analyzed cells refers to Fig. 5f, g.

| **Young's module**<br>**[Pa]**<br>**mean ± standard**<br>**deviation** | **acrylamide, final**<br>**concentration**<br>**[%]** | **bisacrylamide,**<br>**final**<br>**concentration**<br>**[%]** | **analyzed cells**<br>**on Fn** | **analyzed cells**<br>**on Vn** |
| --- | --- | --- | --- | --- |
| 0.4 ± 0.08 | 4 | 0.03 | 44 | 37 |
| 3.0 ± 0.98 | 5.5 | 0.04 | 38 | 52 |
| 6.7 ± 1.95 | 7.46 | 0.04 | 60 | 73 |
| 20.1 ± 4.87 | 7.49 | 0.10 | 54 | 64 |
| 32.6 ± 9.38 | 7.52 | 0.16 | 55 | 50 |
| 61.7 ± 27.03 | 12 | 0.15 | 58 | 61 |

**Supplementary Table 2:** p-values of student t-test (two tailed) for the indicated conditions

| **Young's module**<br>**[Pa]**<br>**mean ± standard**<br>**deviation** | **Fn vs. Vn, cell**<br>**area** | **Fn vs. Vn,**<br>**adhesion length** |
| --- | --- | --- |
| 0.4 ± 0.08 | 0.59 | 0.53 |
| 3.0 ± 0.98 | 0.57 | 0.95 |
| 6.7 ± 1.95 | 0.09 | 0.02 |
| 20.1 ± 4.87 | 0.69 | 0.39 |
| 32.6 ± 9.38 | 0.51 | 0.92 |
| 61.7 ± 27.03 | 0.57 | 0.76 |

## Methods

### Cell culture, constructs, and transfection

NIH 3T3 cells used in this study are a subclone of NIH3T3 cells that were FACS sorted for low expression of endogenous β3-integrin as described previously (Pinon et al., 2014). Vinculin-knockout mouse embryonic fibroblasts (MEF Vcl -/-) and MEF wt were kindly provided by W. H. Ziegler (Mierke et al., 2010). GD25wt cells were kindly provided by R. Fässler (Wennerberg et al., 1996). All cells were grown at 37°C with 5% CO2 in DMEM (ThermoFischer) supplemented with 10% FCS (Hyclone), and passaged 2-3 times a week, or upon reaching confluency. Transfections were carried out with Lipofectamine 2000 (ThermoFischer) or JetPEI (Polyplus) according to manufacturer’s instructions. 6 h after transfection, cells were cultured in complete medium for 48 h before detachment. cDNA encoding full-length mouse β3-wt GFP integrin expressed in a cytomegalovirus promoter-driven pcDNA3/EGFP vector has been previously described (Ballestrem et al., 2001). β3-VE GFP (Pinon et al., 2014), β3-D723A GFP (Ballestrem et al., 2001), and β3-N305T GFP (Ballestrem et al., 2001) were derived by substitution from the β3-wt GFP integrin construct mentioned before and as described in the indicated publications. Vinculin mCherry was a gift from Christoph Ballestrem (Manchester, UK), β5-wt GFP was a gift from Guillaume Montagnac (Gustave Roussy Institute, Inserm), and mApple-MyosinIIA-C-18 was obtained from Michael Davidson (Addgene plasmid # 54929).

### Antibodies and chemicals

Inhibition experiments were performed with blebbistatin (Sigma-Aldrich), with the ROCK inhibitor Y27632 (Sigma-Aldrich), or with the αVβ3 integrin inhibitor cilengitide (Sellekchem) at concentrations as indicated. Dithiothreiol (DTT, Carl Roth) was used at the indicated concentration to open disulphide bridges. Cells were fixed for subsequent immunostaining with 4% PFA (Sigma-Aldrich) in PBS. Reagents used for immunostaining were monoclonal mouse antibodies for paxillin (clone 349/Paxillin, BD Biosciences, # 610052), talin (clone 8d4, Sigma-Aldrich, #T3287), vinculin (clone hVIN-1, abcam, #ab11194), vitronectin (clone VIT-2, IgM, Sigma-Aldrich, #V7881) or polyclonal rabbit antibodies for HA-tag (Sigma-Aldrich, #H6903), fibronectin (Sigma-Aldrich, #F3648), thrombospondin (abcam, #ab85762) or osteopontin (GeneTex, #GTX37582). β1-integrin was stained with a monoclonal rat antibody (clone 9EG7, BD Biosciences, # 553715). After primary antibody staining, samples were washed and incubated with antibodies against mouse labeled with Cy3 (Jackson Immunoresearch, #115-165-146), against rabbit labeled with Alexa Fluor 488 (ThermoFischer, #A11070) or Cy3 (Dianova, #111-165-144), or with phalloidin coupled to Alexa Fluor 568 (ThermoFischer, #A12380). To visualize anti-Vn staining, secondary antibodies against IgM labeled with Cy3 were used (Dianova, #115-166-075). Primary rat antibodies were visualized with preadsorbed, Alexa Fluor 488 or Alexa Fluor 568 labeled secondary antibodies (ThermoFischer, #A11006 or #A11077) and, if present in the experiment, primary mouse antibodies were visualized with preadsorbed, Cy3 labeled antibodies (Dianova, #111-165-144) to avoid cross-reactivity of secondary antibodies. Direct labeling of Fn, Fbg, and Vn was performed according to manufacturer’s protocol with Alexa Fluor 568 (ThermoFischer, # A10238) or Alexa Fluor 647 (ThermoFischer, #A20173).

### Microcontact printing

Silicone stamps for microcontact printing of differential substrates were produced as previously described (Lehnert et al., 2004). Binary choice susbtrates were produced with human plasma fibronectin (Sigma-Aldrich, #F2006 or Millipore, #FC010), human plasma vitronectin (Sigma-Aldrich, #V8379), recombinant human vitronectin (Sigma-Aldrich, #SRP3186), native human fibrinogen (Bio-Rad, #4440-8604), recombinant human thrombospondin-1 (R&D systems, #3074-TH-050), or with osteopontin from bovine milk (Sigma-Aldrich, #O3514). Silicone stamps were incubated for 10 min with a solution of 5 μg/ml Fn and 45 μg/ml heat-inactivated Fn in PBS (see Supplementary Fig. 1i). Alexa Fluor 647 labeled Fn was added to visualize the Fn pattern. After nitrogen drying, the stamp was pressed onto a glass cover slip for 10 min before it was released. Next, the pattern was covered with Vn at a concentration of 1-5 μg/ml in PBS for 1 hour at room temperature. Heat-inactivated Fn was produced by heating Fn for 30 min to 90°C and was added to block Vn adsorption in the stamped area. Cells do not spread on heat-inactivated Fn (Supplementary Fig. 1i). For other binary choice substrates than Fn/Vn, a total concentration of 50 μg/ml of the stamped protein, and 5 μg/ml of the protein for backfilling was used except for Vn/Fn stripe patterns (10 μg/ml for both proteins). All other steps were performed as described. After the final incubation step, patterns were washed with PBS and used directly for cell seeding. Cell detachment from culture flasks was stopped with trypsin inhibitor (Sigma-Aldrich) and cells were cultured for 6 h in DMEM without FCS.

### Polyacrylamide gels

Established protocols (Kandow et al., 2007, Pinon et al., 2014, Plotnikov et al., 2014) were adapted to gain polyacrylamide gels of different Young’s modulus (stiffness) with homogeneous or with structured ECM. Gels were produced on activated cover slips. Therefore, glass cover slips were cleaned first with propanol and second for 10 min in a plasma cleaner (Technics Plasma GmbH, Germany). This was followed by a silanization (1 h at room temperature, 1 mM 3-(Trimethoxysilyl)propyl methacrylate (Sigma-Aldrich) in toluene). After incubation, cover slips were washed in ddH2O and dried with nitrogen. On these cover slips, 60 μl of a mixture of degassed acrylamide, bisacrylamide (both Bio-Rad), tetramethylethylenediamine, and ammonium persulfate (TEMED and APS; Sigma-Aldrich) was pipetted with final concentrations of 0.5% APS, 0.1% TEMED, and as mentioned in Supplementary Table 1. This solution was covered with 10 μl of 1% w/v of Acrylic acid N-hydroxysuccinimide ester (NHS acrylate; Sigma-Aldrich) in toluene. Finally, the solution was covered with a cover slip of 18 mm diameter that was either functionalized with a Fn/Vn pattern prepared as described before or that was coated with a 50 μg/ml solution of Fn or Vn for 1h at room temperature. This top cover slip was dried with nitrogen before it was applied to the gel solution. After polymerization, the top cover slip was removed and the gel was covered with PBS. Gel substrates were used directly for cell seeding or were stored overnight at 4°C before cell seeding. Cells were cultured for 6 h on gels in DMEM without FCS to prevent adsorption of plasma-Vn to the gel surface.

Stiffness of polyacrylamide gels was measured as previously described (Elosegui-Artola et al., 2016). Measurements were performed with the atomic force microscope described below. Silicon nitride pyramidal tips with a nominal spring constant of k = 0.01-0.03 Nm^-1^ were used (MLCT, Bruker). An effective half-angle of 20° was used for calculation. For each stiffness, 3 gels from 3 independent batches were measured by probing 5 positions in the center of the gel with 5 repetitive measurements. The Hertz model equation for pyramidal tips was fitted to the force-displacement curves.

### Microscopy

SR-SIM imaging was performed on a non-serial Zeiss Elyra PS.1 microscope with a 63x/1.4NA oil immersion objective and an Andor iXon EMCCD camera. The grid for SR-SIM was rotated three times and shifted five times leading to 15 frames raw data out of which a final SR-SIM image was calculated with the structured illumination package of ZEN software (Zeiss). Values for calculation were selected for best resolution without causing image artifacts. Channels were aligned by using a correction file that was generated by measuring channel misalignment of fluorescent tetraspecs (ThermoFischer, #T7280). All diffraction limited images according to the figure legend were taken using the ApoTome module on a Zeiss AxioimagerZ1 microscope to achieve optical sectioning. A 63x/1.4NA oil immersion objective and a Zeiss AxioCam MRm were used. For SR-SIM live cell microscopy, the incubation chamber was heated to 37°C and cells were imaged every minute. During imaging, cells were cultured in imaging medium (F12 + 25 mM HEPES + 200 mM L-glutamine + 1% penicillin/streptomycin, pH 7.2). FCS was present as indicated in the description of the Supplementary Movies. SIM raw data images were processed as described above. Phase contrast live cell imaging was performed on a Zeiss Axio-Observer Z.1 with a 20x/0.8NA air objective. Cell migration of GD25 cells on homogenous Fn or Vn (coating: 10 μg/ml in PBS for 1h at RT) was analyzed for cells cultured in DMEM/F12 medium (ThermoFischer, #11039-021) + 1% penicillin/streptomycin + 1% FCS. Migration of GD25 cells on Vn/Fn stripes was analyzed for cells cultured in DMEM/F12 medium + 1% penicillin/streptomycin.

### FRAP

Fluorescence Recovery After Photobleaching (FRAP) was performed as described (Wehrle-Haller, 2007). Image acquisition and image analysis were performed at the Bioimaging Core Facility, Faculty of Medicine, University of Geneva. Briefly, transfected NIH 3T3 cells were cultured on serum coated coverslips. 1 h before imaging medium was replaced with F12 medium (Sigma-Aldrich) containing 10 % FCS + 1% penicillin/streptomycin and cells were relocated to the microscope. FRAP was performed on a Nikon A1r confocal laser scanning microscope equipped with a 60x oil immersion objective and a 37°C incubation chamber. Three pictures in 5 sec intervals were acquired before bleaching. After that we acquired 1 frame / 5 sec for 3 min. The graph was calculated in the following way: The first three images before bleaching were averaged to yield “100% intensity” and the first image after bleaching was set to “0% intensity”. All other values were calculated as ratio of 100% intensity.

### Atomic Force Microscopy

To prepare adhesion substrates for directly comparative adhesion force spectroscopy, adjacent areas on a Fluorodish 35 (WPI) glass bottom dish were coated with 50 μg/ml BSA (to provide low adhesion for cell capture, see below), 50 μg/ml Fn, or 5 μg/ml Vn solutions and incubated for 1h. Substrates were subsequently rinsed five times with PBS and transferred to CO2-independent Medium (ThermoFischer). Prior to SCFS experiments, GD25wt cells were transferred to CO2-independent Medium for 1 h and then trypsinized. Trypsin was subsequently inactivated by adding soybean trypsin inhibitor (Sigma-Aldrich). After centrifugation, the supernatant was removed, and cells were again resuspended in CO2-independent Medium. SCFS experiments were performed using a CellHesion 200 atomic force microscope (JPK) featuring an extended vertical range of 100 μm. All measurements were performed at 37°C using a temperature-controlled sample chamber (BioCell from JPK) and tipless 205 μm long V-shaped cantilevers with a nominal spring constant of 0.06 N/m (NP-O from Veeco). To facilitate cell capture, plasma-cleaned cantilevers were functionalized with concanavalin A. After calibrating the sensitivity of the optical lever system and determining the spring constant, cells were pipetted into the sample chamber. A single cell was captured above the BSA coated area by pressing the functionalized cantilever onto the cell with a contact force of 500 pN for 3 s and elevating the cantilever subsequently. To measure cell detachment forces, the cantilever was lowered at a constant speed of 5 μm/s until the cell made contact with the substrate and a preset force of 1.5 nN was reached. Afterwards, the cantilever was held at a constant height for the preset contact time until the cantilever was elevated 80 μm above the substrate surface. Each cell was tested alternatingly on Fn and Vn surfaces (typically 10 force cycle repetitions for each contact time) to determine the differential adhesion strength to both ligands. In total, 8 different cells were tested. Detachment forces were analyzed using the JPK image processing software. From the collected force-distance curves, the maximum detachment forces (maximum cantilever deflection) were determined and plotted as mean ± SD using OriginPro 8.1G. Statistical significance of experiments was tested with a Wilcoxon-based Mann-Whitney U-test using InStat.

### Image Analysis

Colocalization, cell area, and adhesion length were analyzed with the Fiji software package (Schindelin et al., 2015). A threshold was applied to the intensity of the corresponding fluorescent channel and the area or the length of individual integrin-mediated adhesions was measured with plugins included in Fiji. If necessary, background was subtracted (sliding paraboloid) or analysis was limited to adhesions in areas with less background. Colocalization between two fluorescent channels was quantified by measuring Mander’s coefficient of thresholded images by using the Fiji plugin JACoP (Bolte and Cordelieres, 2006). The location of adhesion initiation on Fn/Vn substrates was defined by analyzing SR-SIM live cell movies. The fluorescent channel of the integrin staining was analyzed while the Fn channel was hidden. Integrin clusters visible for at least two subsequent time frames were marked with an ellipse in the ZEN imaging software throughout the movie. Afterwards, the Fn channel was uncovered, and the positions of all ellipses were counted with respect to FN squares or Vn surrounding the squares. If an integrin cluster initiated at the border of a square with contact to Fn and Vn, it was counted for the category Fn/Vn.

### Biolayer interferometry measurement

Dissociation constants of αVβ3 integrin binding to different ECM proteins were measured using the BLItz biolayer interferometer (Pall ForteBio). All steps during real-time measurements were performed at room temperature in the same buffer conditions (20 mM Tris, 150 mM NaCl, pH 7.4, 1mM MgCl2, 1 mM CaCl2, 0.02% Tween20, 0.1% BSA). Pre-hydrated (10 minutes in integrin buffer) Ni-NTA biosensors (Pall ForteBio) were loaded with 50 μg/ml His-tagged human recombinant αVβ3 integrin (RnD Systems, #3050-AV) following an association phase with 150 μg/ml ECM protein and a dissociation phase (baseline – 45s; loading – 180s; baseline – 45s; association – 300s, dissociation – 300s). Binding curves were corrected for a reference sample, and K_D_ values were calculated from association rate constants, and dissociation rate constants derived by software from local curve fits corrected for start of association and dissociation. Please note that association rate and dissociation rate constants are calculated from the gradient of the respective curve. In control experiments, 20 μM cilengitide was added to the integrin buffer during dissociation phase (Supplementary Fig. 3j). A 10 s adjustment step was included.

### MD simulations

In simulations, we used the structure of FnIII10–bound integrin in extended-closed and extended-open form including propeller, βA, and hybrid domains. Crystal structure of αVβ3 integrin from RCSB Protein Data Bank (PDB: 4MMX) was used as a model for the extended-closed integrin. Structure of the extended-open form was prepared by superimposition of βA and hybrid domains from the crystal structure of open αIIbβ3 integrin (PDB: 3FCU or PDB: 3ZE2 as indicated in the figure legend). Systems containing unliganded αVβ3 were prepared by removing FnIII10 from initial structure. MD simulations were performed using Gromacs ver 2016.1 (Van Der Spoel et al., 2005) at the Sisu supercomputer, CSC, Finland. The CHARMM27 force field (Bjelkmar et al., 2010) and explicit TIP3P water model (Jorgensen and Madura, 1983) in 0.15 M KCL neutral solution were used. Energy minimization of the system was performed in 25 000 steps using steepest descent algorithm. The system was equilibrated in three phases using harmonic position restraints on all heavy atoms of protein. The first phase of equilibration was performed with NVT ensemble for 100 ps using the Berendsen weak coupling algorithm (Berendsen et al., 1984) to control the temperature of the system at 100 K. Integration time step of 2 fs was used in all the simulations. Following NVT, the system is linearly heated from 100 to 310 K over 1 ns using an NPT ensemble at 1 atm of pressure. During this process, the Berendsen algorithm was used to control both temperature and pressure. For the final phase of equilibration and for all subsequent simulations, an NPT ensemble was maintained at 310 K, using V-rescale algorithm (Bussi et al., 2007), and 1 atm as implemented in Gromacs 2016.1. Temperature coupling was applied separately for protein and solution parts. Preparation of structures and analysis was performed using PyMOL 1.7. Interaction energy, which was calculated to evaluate RGD-integrin interactions, is a sum of Lennard-Jones and Coulomb energies.

### Statistics

If not stated otherwise, reported values in bar charts are calculated as mean and error bars are representing standard deviation of all data points. In box plots, upper and lower bar indicate standard deviation and the middle bar indicates the mean. Statistical comparisons are calculated with two-tailed Student’s t-test based on the number of independent experiments. For adhesion force measured with AFM, statistical significance of experiments was tested with a Wilcoxon-based Mann-Whitney U-test using InStat. All experiments were reproducible and were carried out as independent experiments at least twice or as often as indicated in the figure legends.

## List of abbreviations

Fbg: fibrinogen
Fn: fibronectin
FnIII10: 10^th^ domain of fibronectin
MEF: mouse embryonic fibroblast
MEF vcl -/-: mouse embryonic fibroblasts derived from vinculin knockout mice
MD: molecular dynamics
NMIIA: nonmuscle myosin IIA
Opn: osteopontin
Tsp: thrombospondin
Vn: vitronectin

